# Modular integration of neural connectomics, dynamics and biomechanics for identification of behavioral sensorimotor pathways in *Caenorhabditis elegans*

**DOI:** 10.1101/724328

**Authors:** Jimin Kim, Jeremy T. Florman, Julia A. Santos, Mark J. Alkema, Eli Shlizerman

## Abstract

Computational approaches which emulate *in-vivo* nervous system are needed to investigate mechanisms of the brain to orchestrate behavior. Such approaches must integrate a series of biophysical models encompassing the nervous system, muscles, biomechanics to allow observing the system in its entirety while supporting incorporations of different model variations. Here we develop *modWorm*: a modeling framework for the nematode *Caenorhabditis elegans* using *modular integration* approach. modWorm allows for construction of a model as an integrated series of configurable, exchangeable *modules* each describing specific biophysical processes across different modalities (e.g., nervous system, muscles, body). Utilizing modWorm, we propose a base neuro-mechanical model for *C. elegans* built upon the complete *connectome.* The model integrates a series of 7 modules: i) intra-cellular dynamics, ii) electrical and iii) chemical extra-cellular neural dynamics, iv) translation of neural activity to muscle calcium dynamics, v) muscle calcium dynamics to muscle forces, vi) muscle forces to body postures and vii) proprioceptive feedback. We validate the base model by *in-silico* injection of constant currents into sensory and inter-neurons known to be associated with locomotion behaviors and by applying external forces to the body. Applications of *in-silico* neural stimuli experimentally known to modulate locomotion show that the model can recapitulate natural behavioral responses such as forward and backward locomotion as well as mid-locomotion stimuli induced responses such as avoidance and turns. Furthermore, through *in-silico* ablation surveys, the model can infer novel neural circuits involved in sensorimotor behaviors. To further dissect mechanisms of locomotion, we utilize modWorm to introduce empirical based variations of intra and extra-cellular dynamics as well as model optimizations on associated parameters to elucidate their effects on simulated locomotion dynamics compared to experimental findings. Our results show that the proposed framework can be utilized to identify neural circuits which control, mediate and generate natural behavior.

## 1. Introduction

Neural circuits within the nervous system use rhythmic activity to facilitate coordinated body movements. Central Pattern Generator (CPG) networks are thought to generate rhythmic neural activity and motor behavior in organisms such as the locust, lamprey, *Drosophila* and the nematode *C. elegans* (1–7). However, CPG networks alone do not explain many details of sensorimotor integration and the functional pathways guiding neural activity and movement. Computational approaches which integrate the full nervous system are thus needed to identify neural circuit candidates mediating behavior. An iterative approach of discovering sensorimotor pathways via computation followed by empirical validation has the potential to discern the fundamental principles through which the nervous system and body interact (8).

In this respect, it is appealing to study the nematode organism *C. elegans,* in which inherent locomotion patterns are well observed and characterized (9). These along with environmental stimuli which lead to a change of locomotion direction provide an intriguing and well quantified model organism for locomotion investigation. *C. elegans* neuronal wiring diagram, which maps electrical and chemical neural connections between somatic neurons within its nervous system is resolved and constantly being updated across organism’s sexes and developmental stages (10–20). The availability of connectome warrants searching for incorporated circuits using computational and experimental techniques. Indeed, groups of sensory-, inter- and motor-neurons have been associated with various types of locomotion including natural crawling motions (21–23), chemosensation (24–30), thermosesation (31–33) and mechanosensation (34–37). It is still not fully resolved, however, how these sensorimotor mechanisms are incorporated on a network level in *C. elegans* and what types of neural interactions lead to locomotion behaviors (38–40).

A central reason for the complexity stems from biophysical dynamics additional to the connectome. These dynamics encompass the processes representing neural responses and body bio-mechanics (22,41–46). Such processes add numerous and intricate possibilities for neural signals to flow within the routes set by the connectome. Indeed, sensorimotor integration within *C. elegans* neuron’s individual responses is likely to be highly recurrent and interactive through synapses, gap junctions, neuromodulators, extra-synaptic signaling, muscles, body, proprioception, etc (46–56). While it appears as a complex system to study, the fact that these processes are coordinated during locomotion suggests that their compound effects can be inferred through the integration of individual models that emulate these processes (48,57,58). Integrating such models with varying scopes and modality warrants the development of an effective modeling framework that allows for both simultaneous simulations and testing of individual models. We therefore develop a framework allowing such a modeling approach and refer to it as *modWorm*: a modular integration modeling framework for *C. elegans* neuro-mechanics. The framework allows construction of a model as a series of configurable, exchangeable biophysical *modules* each responsible for modeling distinct dynamic processes (as depicted in Fig. 1B).

**Figure 1:**
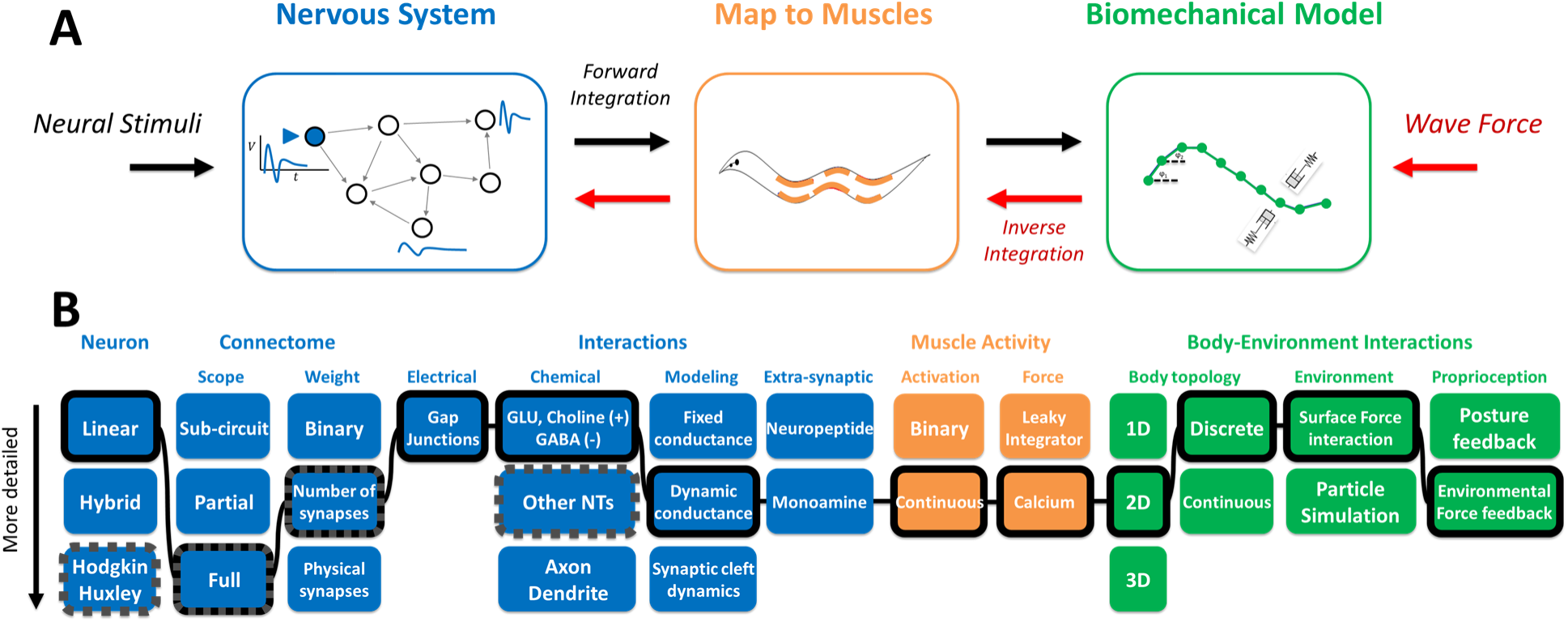
Constructing *C. elegans* neuro-mechanical model. **A:** From left to right, Modeling the nervous systems as a dynamical system encompassing the full somatic connectome including graded ion-channel and connectivity neural dynamics, mapping neural dynamics to dynamic muscle impulses and forces, mapping muscle forces to a biomechanical model that incorporates body responses and interaction with the environment. Neural stimuli are integrated forward to resolve body movements (black arrow). External forces are propagated in an inverse direction to resolve corresponding neural dynamics (red arrow). **B:** From left to right: Schematics of each model aspect as flowchart for the nervous system (blue), map to muscles (orange) and biomechanical model (green). Black highlighted boxes connected with solid lines are model aspects chosen by the proposed base model and boxes with dark gray dotted edges represent the variations considered in the paper.

Finding a combination of modules which constitute locomotory responses involves considering survey of a large pool of possible model variations. In the context of modeling *C. elegans* neuro-mechanics, this amounts to more than 50,000 variations over 13 different model aspects encompassing the nervous system, muscles and biomechanics, each with varying degrees of complexity (Fig 1B). Since it would be computationally infeasible to fully survey such large number of variations, we take a more fundamental approach where we construct an initial model comprised of *base* aspects that are generic, scalable, and unbiased by specific behaviors or experiments (i.e., no parameter fitting). These model aspects are chosen based on experimentally found anatomical and electrophysiological data as well as biophysical processes that exist in *C. elegans* (Fig 1B) (59–66). Using modWorm, the base model is constructed with a total of 7 modules incorporating simulation of the nervous system as a dynamical network, translation of neural activity to muscle forces, and mapping muscle dynamics to *C. elegans* body model.

The proposed base model of *C. elegans* integrates the full known somatic connectome with linear intracellular and non-linear extracellular neural dynamics driven by detailed synaptic transmission model similar to reduced Hodgkin-Huxley model type ion channel. The model further integrates body biomechanics and its interactions with the environment by translating the activity of the nervous system to calcium driven muscle forces which in turn generate body postures according to surrounding environment-body interactions. In addition, it implements proprioception in the form of inverse integration which modulates neural dynamics according to external forces on the body. We test and validate these model aspects by analyzing simulated neural dynamics driven by both external wave forces and neural stimuli associated with locomotion as well as studying the effects of environmental variations and proprioceptive feedback to simulated body dynamics. We also extend our studies to incorporate dynamic stimulus to investigate complex behaviors such as avoidance and turns induced by timed stimulus.

The base model permits in-place modifications to any of its modules to study its effects on neural and body dynamics. We therefore use this capability to perform *in-silico* ablations on neurons or connections and infer neural circuits involved in sensorimotor behavior associated with a stimulus. Implementation of such approaches for the study of turn response triggered by stimulus shows that we can identify neural pathways that facilitate such behavior. We use systemic ablation to also recapitulate locomotion behaviors associated with *in-vivo* ablation experiments of touch responses and perform further ablations to elucidate novel details on these experiments.

The modular structure of the base model also allows for its underlying modules to be extended or exchanged to test the refined fit of locomotion to empirical findings. We thus introduce different variations to the base model ranging from individual neuron channels, synapses to full connectome mappings to study their effects on simulated locomotion and identify variation which results in the most significant improvement in simulation quality. In addition to variations that are empirically based, we also consider model optimization through neural parameters fitting (e.g., synapse strengths) with respect to a given locomotion task and an alternative module for extra-cellular neural dynamics to elucidate their effects on simulated behavior. Testing such variations highlights the possible mechanisms of the investigated locomotion and directions in which the base model can be improved and how it can be incorporated into *in-vivo* investigations.

## 2. Related works

Models of neural dynamics and biomechanics have been introduced for several model organisms including adult and larval *Drosophila* (20,67–72), hydra (73), lamprey (74,75), leech (76,77) and rodents (78). For *C. elegans*, the availability of anatomical, electrophysiological, biomechanical, behavioral data makes the organism a suitable candidate for neuro-mechanical modeling (12,13,16,18,79–81). Indeed, several models incorporating neural and body dynamics of varying scopes have been proposed. Here we survey these models categorized into several broad approaches to highlight their contributions and differences with respect to modWorm.

### Models incorporating partial connectomes

Several models integrating ***partial connectome data*** (e.g., sub neural circuits) with potentially biomechanics models have been introduced. Such models include methods describing ventral motor neurons as symmetric binary units which control the body of *C. elegans* segmented as discrete rods and stretch receptors (82). The model showed gaits generating forward locomotion but also locomotion instabilities when neurons dynamic properties and arrangement are slightly changed, e.g., when binary motor units are replaced by neural dynamics. Several studies also showed that pattern generators or connectome based locomotory sub-circuits (e.g., head motor neurons and ventral nerve cord) combined with body models can produce forward locomotion and basic navigation behavior such as Klinotaxis (24,83–90). While these analyses show that oscillators and sub neural circuits can produce *C. elegans* forward locomotion body postures and subsequent sensory navigation, their relation to the full nervous system and how these patterns are being generated remains unresolved. Furthermore, a unifying relation to other locomotion behaviors such as backward, turns and pirouettes remains unclear.

### Models incorporating complete connectomes

Other lines of connectome based models introduced a dynamical model for the complete somatic nervous system (91–94). These studies showed that the full nervous system can generate dynamic rhythms even when a few mechano-sensory neurons received a constant stimulus. These rhythms, however, could not be directly associated with behaviors since additional processes of biomechanics and proprioception were not included. Inspired by the human brain project, the OpenWorm collaborative project was established in 2011 as a crowdsourcing platform aimed to develop generic bottom-up simulations of neuronal models, body and fluid simulations to lead to a full-scale *C. elegans* model (15,91,95). While there has been progress in the development of generic tools for modeling *C. elegans* and other organisms, such as Geppetto, c302 (multiscale modeling) and Sibernetic (hydrodynamic simulation) (95,96), integration of these tools into an unified framework has not yet been achieved. Moreover, incorporation of feedback between the nervous system and body such as proprioception remains unresolved.

### Inclusion of ML driven models for enhancement

Recent works include methods inspired by Zador et al (97) adopting machine learning techniques to enhance the model and improve its simulation accuracy. These methods target parameters such as neuron polarities, connection weights/strengths and muscle-body parameters to be optimized using machine learning algorithms with respect to the established empirical data. Such methods have been applied to nervous system of both *C. elegans* and *Drosophila* to infer particular neural functions (98–101). Similar approaches have also been developed for the neuro-mechanical modeling of *C. elegans* to reproduce stereotypical behaviors such as forward locomotion (102,103). These models, however, generally include a number of compromises on model details, such as using partial connectome, less accurate first order approximation for neural dynamics, and absence of proprioception or body-environment interactions, to accommodate large scale optimization algorithms (e.g., Back-propagation through time) (104). Furthermore, their ability to simulate locomotion behaviors additional to forward locomotion such as backward, turns and pirouettes, etc are not guaranteed.

Here, we introduce a modular modeling framework: modWorm, and subsequent neuro-mechanical model of *C. elegans* that aims to address these limitations via simultaneous simulations and observations of the system in its entirety. The model integrates established biophysical processes and their approximate parameter values in *C. elegans* which encompass the complete somatic nervous system, muscles, body, and their interactions with the environment. We show that such a base model incorporates and qualitatively reproduces known forward and backward locomotion, and transitional behaviors such as turns and avoidance. By introducing experimentally driven variations and optimizations to the base model (e.g. parameter fitting), we further show the effects of each variation on simulated neural and body dynamics with respect to their base counterparts and experiments. Such an approach of implementing initial base model, followed by its variations and experimental validations can assist in making informed decisions in which the model incorporates and reflects future details and *in-vivo* findings into an expanding model of the organism (8,105–107).

## 3. Results

### 3.1. Base *C. elegans* neuro-mechanical model

We utilize modWorm to construct *the C. elegans* neuro-mechanical model. The model consists of 3 constituent models (*Nervous System, Muscles, Biomechanics*) comprised of 7 modules (Fig S1). The Nervous System Model adopts recently established dynamical model of *C. elegans* neuronal network which simulates responses of the full somatic nervous system (279 neurons) to stimuli (83,92,94). The model is based on molecular properties of neurons in *C. elegans* network and describes neural responses as combination of 3 modules: (i) graded potentials (ii) neural gap-junction connectivity (iii) neural dynamic synaptic connectivity including glutamatergic, cholinergic and GABAergic receptors. In the base model, glutamatergic and cholinergic transmitter activated ion channels are assumed to be excitatory, and GABAergic receptors are assumed to be inhibitory. These settings are configurable to changes (as we show in the section *Model variations for investigation of simulated behaviors*). The synaptic dynamics are modeled as Hodgkin-Huxley model type ion channels where the transmitter release is controlled with dynamic gating variable that is simulated alongside the membrane potential. The model incorporates local synaptic parameters determined by the connectome (i.e., number of synapses) and global biophysical conductance coefficients per type of connection. For more details on the Nervous System Model, see SM (Supplementary Materials) and (13,14).

The Nervous System Model allows for computational clamping of neurons by external current injection. It was previously observed that injection of constant current into model’s sensory neurons, e.g., the posterior PLM mechanosensory neurons, evokes oscillatory neural responses in subset of motor neurons producing low dimensional attractor-like dynamics and transient dynamics with longer timescales than the intrinsic neural dynamics (93,108). Since our initial goal is to implement a base model that is generic and not specific to behavior or experiment biases, we do not fit the neural parameters of individual neuron channels and synapses. These values, however, are easily configurable through modWorm. In the section *Model variations for investigation of simulated behaviors* and Fig. 7, we perform plausible modulations to such parameters to examine the effect of additional experimental details or hypotheses (16,109–112).

We integrate the Nervous System Model with Muscles and Biomechanics models for the investigation of how the nervous system transforms its dynamics to behaviors (Fig. S1). Muscles and Biomechanics models consist of 3 modules describing (i) translation of the nervous system activity to muscle calcium dynamics, (ii) calcium dynamics to muscle forces, (iii) muscle forces to body postures according to surrounding environment-body interactions. The Muscles Model connects motor neurons excitations to muscles activations using an experimentally determined linear map (15,81) (Fig. S6). The Biomechanics model then translates muscle activations to body dynamics by resolving the interactions between the musculature of *C. elegans* (modeled as two-dimensional viscoelastic rod) and external force from surrounding fluid environment. Such an approach was proposed for investigation of eel swimming with muscles activated by external signals emulating neural excitation (113,114). We implement this model rather than the gaited model proposed for *C. elegans* or three-dimensional body model with full particle simulations (15,82,115). This is since body discretization into segments has shown to be stable under dynamic neural stimulation. Moreover, modeling the body as two-dimensional rods and body-fluid interactions in the form of damping are effective methods of simulating *C. elegans* body in a 2D plane and its surrounding environment while maintaining computational efficiency. Indeed, in the section *The effect of the environment on locomotion*, we show that such a model can emulate locomotion body postures under environmental variations similar to *in-vivo*. In previous work, statistical models, partial connectome, or synthetic muscle stimulation were used to generate body movements (82,86,115–117).

The implementation of 2D body model allows us to simulate the complete somatic nervous system simulation (279 neurons) for such excitation. See SM for more detailed descriptions of Muscles and Biomechanics Models.

### 3.2. Locomotion and corresponding neural dynamics induced by external force

In addition to integrating the model in a feed forward manner, we develop an inverse integration approach. Inverse integration computes the neural dynamics that would result from external forces applied to the body. The approach transforms external forces acting on the body to muscle activity and then inverts the activity to membrane potential. These potentials are then integrated forward to resolve the body posture (see SM for more details). We use this approach to validate our model for three basic locomotion patterns: forward, backward and turn movements. For each pattern we apply a sinusoidal force wave travelling along the body with variable frequency to infer neural dynamics associated with it. These neural dynamics are then forward integrated by the nervous system to generate the body posture. We then simulate the integration and compare with the locomotion characteristics of freely moving animals (see SM Videos, snapshots, curvature maps, and calcium activity, deviation in first 6 eigenworm coefficients in Fig. 2).

**Figure 2:**
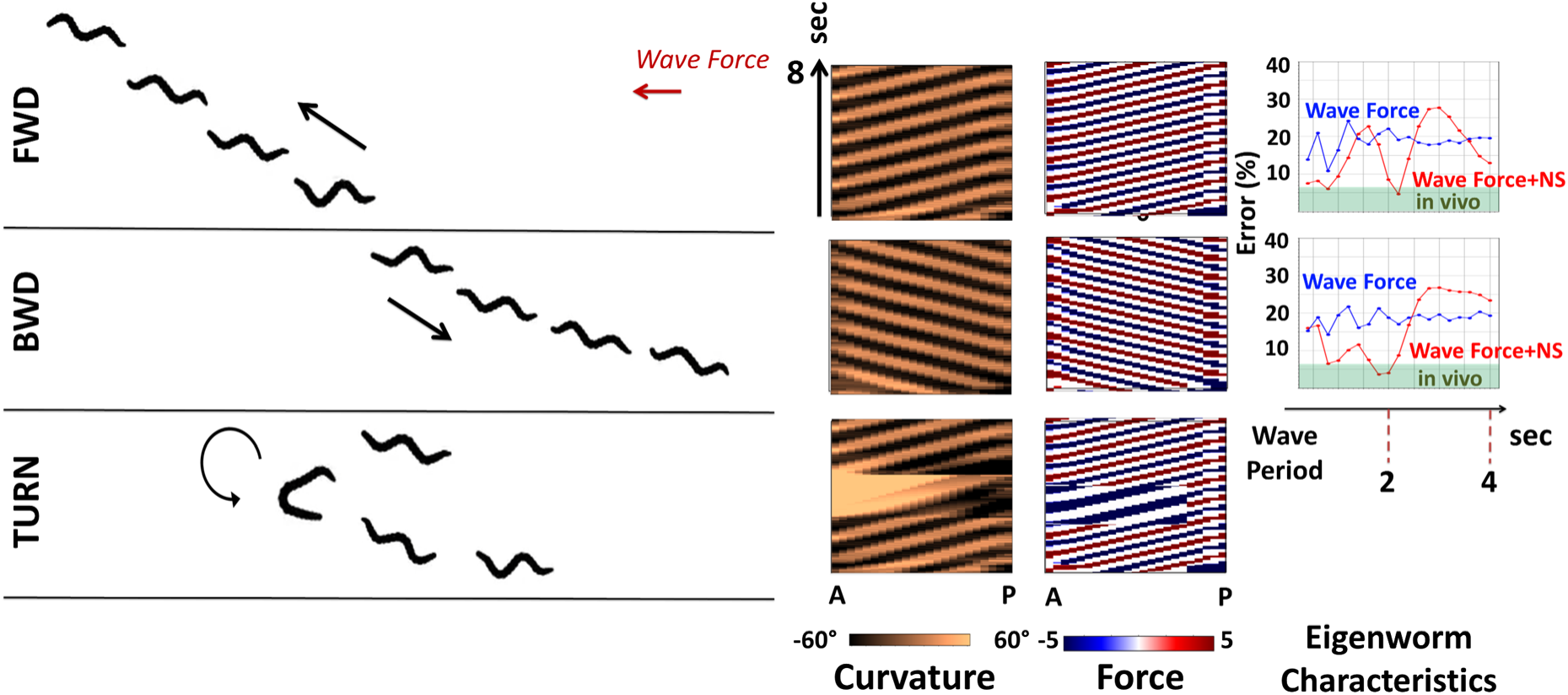
Typical locomotion patterns, body curvature and force dynamics generated by three types of external wave forces, corresponding to forward, backward, 180° turn movements. From left to right: Simulated body snapshots driven by external wave force with minimal eigenworm posture error where each snapshot is sampled every 2 seconds for forward, backward and turn dynamics (Also see SM videos), associated body curvature and muscle force dynamics ordered from anterior to posterior direction for 8 seconds simulation duration, eigenworm posture errors (w.r.t normalized posture coefficients) in the function of varying external wave force periods. Posture errors resulting from direct external wave force on the body are labeled as blue curve with ‘Wave Force’. External wave force inverse integrated to resolve neural dynamics which are integrated forward to simulate body dynamics are labeled as red curve with ‘Wave Force+NS’. ‘Wave Force’ curve results with ∼20% mean error for both directions of the force and does not show preference to wave period. ‘Wave Force+NS’ appears to be selective to period and achieves minimal error for period ∼ 2s of 4.6% (fwd) and 3.6% (bwd) within the CI of *in-vivo* worms (green band; 6.7%, P=0.01) for both directions of movement.

We use eigenworm characteristics to compare the effect of inverse integration of external forces through the nervous system versus external forces acting directly on the body with no nervous system (9,118,119). Postures generated by the integration of the nervous system result in a close match to the coefficients of freely moving worms with preference for particular periods (4.6% and 3.6% normalized coefficient errors within 6.7% *in-vivo* error interval obtained from freely moving animals). The optimal frequency of the force that is being selected is approximately 2s. These results indicate that the response of the nervous system is shaping the external effect on the body in a nontrivial and nonlinear manner (Fig. 2).

Next, we investigate the neural dynamics associated with these locomotion patterns. In Fig. 3A, B we show membrane potential traces for the full somatic nervous system, and a group of 58 motor neurons part of the Ventral Nerve Cord (DB, VB, DA, VA, VD, DD) reported in terms of difference from the rest membrane potential. We observe that the traces are qualitatively consistent with activity patterns identified in the literature (120–123). Most of the neurons are activated during locomotion where (DB, VB) group is the most active group in forward locomotion and (DA, VA) group is the most active group in backward locomotion. When we focus on (DB, VB) neurons and order them by their physical location along the anterior to posterior axis, we find that within each period of oscillation, the activity propagates with a preferred spatial direction (Fig. 3C). During forward locomotion, membrane potential activity propagates from Anterior to Posterior (A→P) while for backward locomotion, the propagation is from Posterior to Anterior (P→A). These propagation directions are consistent with the direction of movement (124,125).

**Figure 3:**
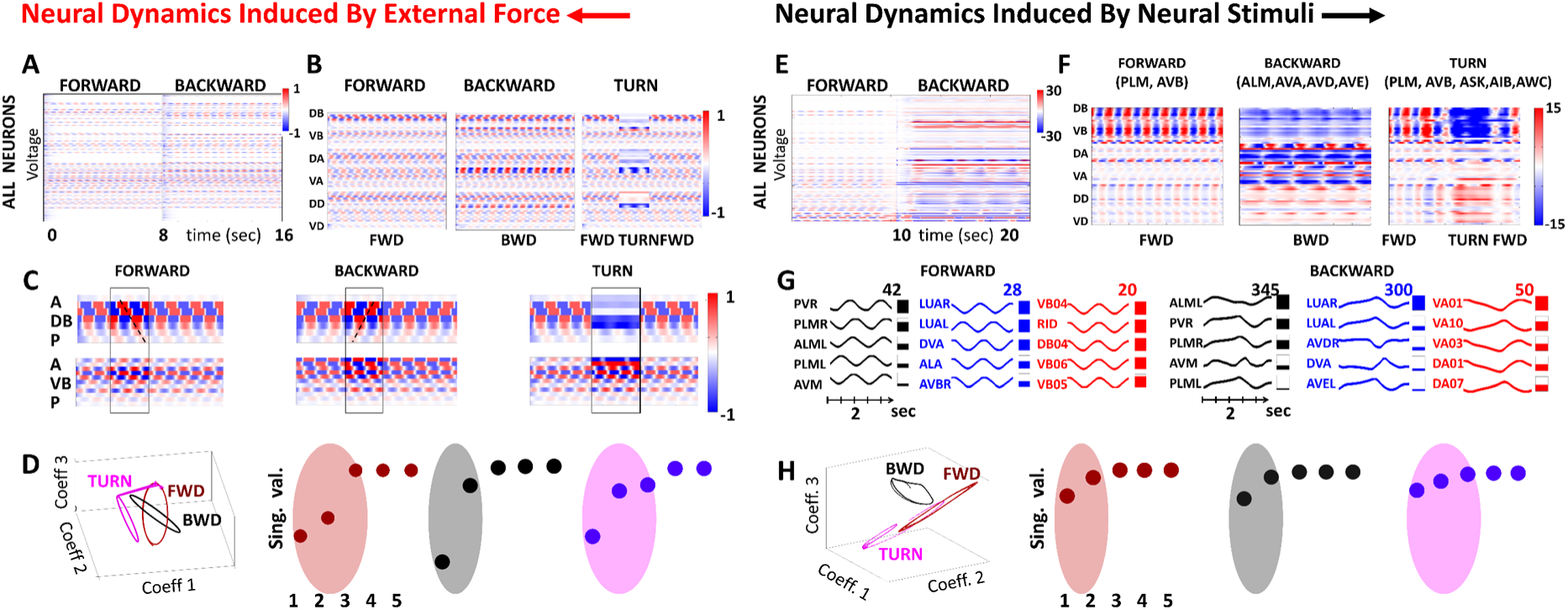
Neural responses of *C. elegans* somatic nervous system to external wave forces and neural constant stimuli. **A**: Color raster plot of membrane potential (difference from equilibrium) of 279 neurons inferred by inverse integration of wave forces to corresponding neural dynamics. Neural responses generated for 16 sec: 0-8 sec: spatial wave force generating forward movement; 8-9: transition; 9-16 sec: spatial wave force generating backward movement. **B**: Color raster plots of membrane potential of motor neurons for forward, backward and turn wave force profiles. **C:** Color raster plots of membrane potential of Ventral and Dorsal type B motor neurons for forward, backward and turn wave force profiles **D**: Evolution of temporal coefficients during forward, backward and turn neural responses (red, black, magenta). Temporal coefficients are associated with PC modes from SVD analysis of all three responses (i.e. projected to a common space of PC1-PC3). **E:** Color raster plot of membrane potential of optimal forward and backward current; 0-10s: forward (0.7nA into PLMR/L, 1.3nA into AVBR/L) 10-15s: transition; 15-25s: backward (2.8nA, 1nA, 0.5nA, 0.5nA into ALMR/L, AVAR/L, AVDR/L and AVER/L respectively). **F:** Color raster plots of membrane potential of motor neurons for forward, backward and turn stimulations (compare with Figure 3B). **G:** Top 5 neurons (which have largest elements in PC1 mode) from each group (sensory, inter and motor) of neurons for forward (left) and backward (right) stimuli. **H**: Evolution of temporal coefficients during forward, backward and turn neural responses (red, black, magenta); compare with Figure 3D.

Analysis of membrane potential responses using Singular Value Decomposition (SVD) elucidates their low dimensional characteristics. The SVD method decomposes the responses into spatial neuronal population modes (PC modes) and their temporal coefficients (92,121,126). We first apply SVD to understand the representation of each individual movement to determine the number of spatial modes needed to represent each activity. The decomposition reveals that there are only a few (2-3) dominant spatial modes representing each movement similar to empirical findings (127). We thus use these modes to construct unified low-dimensional basis of spatial neuronal modes, to elucidate discriminative signatures of forward and backward movement (122). A viable candidate for such a basis is the set of the first three PC modes obtained from SVD of all motor neurons membrane potential during forward, backward and turn movements. Projection of forward and backward responses onto this basis (PC space) yields cyclic temporal trajectories which are well separated and appear to be orthogonal (Fig. 3D). When projecting the turn membrane potential dynamics onto this space we observe that the trajectory departs from the forward cycle and approaches the region of the backward cycle, highlighting the shift in neural dynamics. Notably, the PC space and the coefficients trajectories are obtained from the raw membrane potential dynamics and not from the derivative of calcium dynamics as described previously (122). This allows us to identify the PC space as a low dimensional recognition space capable of determining the type and characteristics of movements the network performs from motor neural activity.

### 3.3. Neural dynamics induced by neural stimuli

In complement to external body forces, we apply neural “clamping” to examine how these stimuli generate body movements. Our first aim is to explore movements created from simple constant stimuli where most of the neurons do not receive any input. In later investigations we extend the stimulus to be a dynamic timed stimulus (Fig. 6). To identify stimulus profile which supports coherent locomotion, we target the circuit associated with the touch response and seek for stimulations of sensory- and inter-neurons maximizing locomotion distance in either forward or backward directions (34,120,128). From neurons in this circuit, our simulations identify a subset of both sensory- and inter-neurons related to behavioral responses: posterior-touch triggered forward locomotion (sensory: PLM (0.7nA), inter: AVB (1.3nA)), anterior touch triggered backward locomotion (sensory: ALM (2.8nA), inter: AVA (1nA), AVD (0.5nA), AVE (0.5nA)), and turn movement (sensory: +ASK (0.3nA), AWC (0.6nA), inter: +AIB (0.5nA)). See Fig. S2 for illustration of optimizing the stimuli amplitudes into these neurons resulting in maximum locomotion distances. The results show that stimulation of both sensory and interneurons promote directed locomotion as reported in experimental studies and control theory analyses (129,130).

Membrane potential traces associated with neural clamping are consistent with membrane potential activity generated by external force traveling waves. We observe similar active groups of motor neurons: (DB, VB) for forward, (DA, VA) for backward, a phase change in (DB, VB) in turn (Fig. 3E), and similar preferred spatial direction in each period of oscillation. SVD analysis on membrane potential traces indicates occurrence of dimension reduction similarly to the external force case. We select the top five neurons from each neural group (sensory, inter, motor), which received the highest weight in the first PC mode, and display their membrane potential over 4 sec time (Fig. 3G). Notably, the selected neurons are those that are experimentally linked to forward and backward movements; for forward locomotion PLM, PVR sensory neurons and VB motor neurons, and for backward ALM, PLM sensory neurons and VA motor neurons. Furthermore, membrane potential response time patterns are characteristic to the two different types of locomotion examined: for forward stimulus these are clear sinusoidal membrane potential dynamics with period of ∼1.8sec in- and out-of-phase oscillations and for backward stimulus these are cusp like responses over longer period of ∼3.4sec with two, in- and out, phases as well. These oscillations are not present in the stimulus and are generated by the intrinsic neuronal network interactions. Projection of forward, backward locomotion and turn responses onto 3 PC modes embedding yields well separated cyclic trajectories as in the spatially traveling wave stimulation: the forward cycle is approximately orthogonal to backward cycle and turn membrane potential projected trajectory departs from the forward cycle to approach the region of the backward cycle (Fig. 3H).

### 3.4. The effect of the environment on locomotion

Experiments indicate that the environment plays an important role in shaping coordinated movement (124,131,132). We thereby explore environmental variations and their influence on the model with respect to parameters such as viscosity of the surrounding fluid and rod elasticity, which represents the ability of the rod to propagate forces along the body (133). In all variations, we fix the neural stimulus to simulate forward locomotion as in Fig. 3 and examine the effective body movement. We observe that as the environment varies, there are changes in the global characteristics of the movement as demonstrated in Fig. S10. For fluid viscosity variation, we compare the postures generated by the model to *in-vivo* experiments in which the viscosity of the surrounding agar fluid was altered by dropping a water droplet (lower viscosity than agar) (134). We mimicked such an experiment by simulating the locomotion in two different viscosity values corresponding to either agar (10mPa/s) or water (1mPa/s). We observe that for the control environment (agar), the model generates movement that features a *sine*-shaped posture qualitatively similar to that of the experiment, whereas for low viscosity (water) the same stimulation corresponds to strokes of C-shaped postures, atypical to *C. elegans* forward motion (Fig. 4A). When compared to experimental body curvatures in water, these postures have similar overall C-shaped postures as demonstrated in Fig. 4A and accompanying SM videos. These results show that a particular neural activity can support various types of behavioral responses, which are determined not only by precise choice of command neurons stimulus, but also by appropriate surrounding media for the movement.

**Figure 4:**
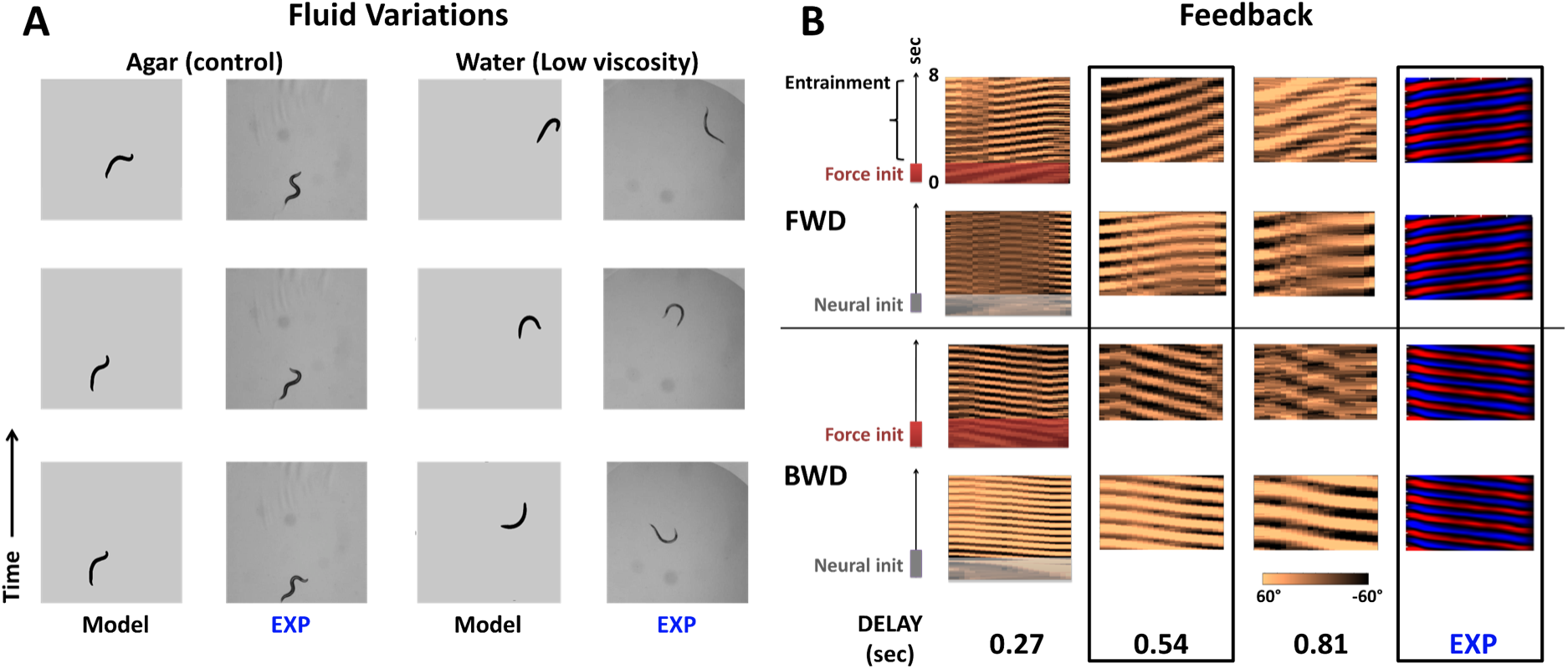
Appropriate fluid parameters and proprioceptive feedback and facilitate sustained locomotion. **A**: Surrounding fluid of model and experiment (134) are varied between agar (high viscosity, columns 1,2) and water (low viscosity, columns 3,4) respectively, to study their effects on locomotion. In the model, viscosity and fluid density are reduced from 10mPa/s, 1g/cm^3^ to 1mPa/s, 0.7g/cm^3^to emulate water droplet in experiment. For each column, 3 locomotion snapshots are taken at 1 second apart. **B**: Feedback is initiated by a wave force (rows 1,3) or neural stimuli (rows 2,4) (see also SM Videos). Columns 1-3 display various feedback time delays, modeling the environmental reaction time of producing external forces on the body, and the curvature profiles they produce and compared to column 4, experimentally recorded curvatures (adapted from (125)). Time delay of approximately 0.5 sec produces optimal forward and backward locomotion which is close to experimental locomotion (highlighted by dashed border).

### 3.6. The effect of feedback (proprioception) on locomotion

Experiments indicate that proprioception within the motor neurons circuit can facilitate locomotion and is an alternative to stimulation of command interneurons (117,131,135–139). To emulate proprioceptive feedback, we incorporate an additional inter-cellular dynamics module into the base model which outputs neural stimulation term I^FDB^ based on external body forces by inverse integrating the force that acts on the body subject to a *time delay* (Fig. S1, see SM for details). Such time-delayed feedback control has been suggested to play important role shaping coherent locomotion for anguilliform swimmers (75,87,140). We then test feedback effects by initiating locomotion with external stimulation, either neural or external force stimulation. Once the feedback starts to entrain the movement, we gradually turn the stimulation off. We found that in both initiation procedures, feedback entrains the body into sustainable coherent movements in forward and backward directions, such that the body moves solely due to feedback (Fig. 4B; SM Videos). Variation in feedback delay time influences coherency and we find the delay of approximately 0.5 sec to be closest to experimentally measured patterns (Fig. 4B) (125).

### 3.7. Case study of the touch responses

#### 3.7.1. Validation, recapitulation and prediction of neural ablation effects

Next, we link neural stimulations with proprioception feedback and examine locomotion responses studied in the literature for (i) *gentle* and (ii) *harsh* anterior/posterior touch. Neural and mechanical triggers for these behaviors have been identified and we use them to validate the movements generated by base model (34,37). For all four touch responses, the base model generates typical directional movement patterns (i) forward; during posterior touch neural stimulation (PLM (Gentle) and PVD+PDE (Harsh)) and (ii) backward; during anterior touch stimulation (ALM+AVM (Gentle) and FLP+ADE+BDU+SDQR (Harsh)), as shown in Fig. 5A (Control (Wild)). To test how robust the recapitulation of the response is, we perform *in-silico* ablations of random pairs of neurons (Fig. 5A; Control (Rand)) as additional controls. Indeed, random ablations do not change, on average, the characteristic velocities of the four touch responses that we considered. Next, we consider *targeted in-silico* ablations as done in prior *in-vivo* experiments and compare velocities and directions of movements with respect to the descriptions published in these experiments (34,37); See Fig. 5A. Arrows indicate *in-vivo* reported data; color of arrows indicates consistency (blue); or disparity (red).

**Figure 5:**
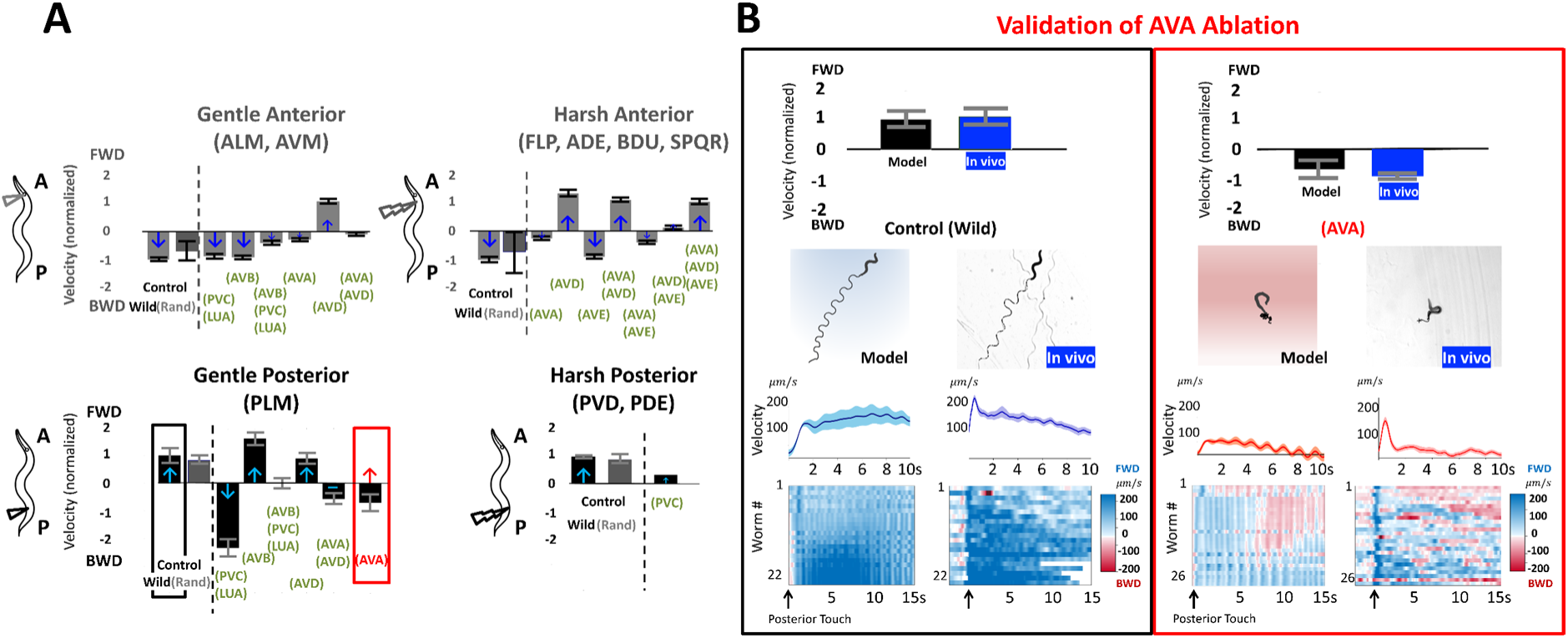
Validation, recapitulation, and prediction of locomotion behaviors for touch responses. **A:** Body velocities for neural stimulation associated with Top: Anterior Gentle (left) and Harsh (right) touch responses Bottom: Posterior Gentle (left) and Harsh (right) touch responses. Velocity amplitude is normalized according to the Control Wild (black label) locomotion and direction up/down are chosen as FWD and BWD direction respectively. Recapitulated velocities following ablation experiments in (34, 37) (green label) are compared to the Control Wild velocities. Arrows indicate experimental *in-vivo* observations and color indicates match of model with published experimental observations (blue: match; red: mismatch) **B:** *In-vivo* validation of Gentle Posterior Touch model prediction for ablation of AVA. Model predictions are compared with *in-vivo* assays for control (left) and AVA ablated (right). Model and *in-vivo* responses at and after stimulus onset are compared with characteristics of (top to bottom) normalized velocities, response locomotion paths, instantaneous velocities after onset, individual instantaneous velocities (each row is a worm) color plot (blue; positive velocity (fwd); red: negative velocity (bwd)).

Our ablation results are in general, qualitatively consistent with previous *in-vivo* findings (19 scenarios out of 20). For the gentle anterior touch responses (left-top in Fig. 6A) ALM, AVM we observe that ablations of (AVA) or (AVD) are the most impactful. Ablation of (AVA) nearly stops movement, while ablation of (AVD) reverses the direction to forward movement in some cases with respect to the control response of backward movement. Notably, effects of *in-silico* ablation are not simple nor additive, e.g., ablation of (AVD) alone is stronger than ablation of (AVA) + (AVD); ablation of (AVB) + (PVC) + (LUA) causes the movement to significantly slow down, while separate ablation of (AVB) or (PVC) + (LUA) permit the typical backward movement. Similarly, for the stimulation (PVD, PDE (Harsh posterior scenario); right bottom in Fig. 5A) we observe that ablation of (PVC) does not lead to a reverse in direction as in *in-vivo* experiment. For (FLP, ADE, BDU, SDQR (Harsh anterior scenario); right-bottom in Fig. 5A) ablation of (AVD) or (AVA) + (AVD) is found to reverse the direction from backward to forward. This observation is similar to the observation described in *in-vivo* experiments (34,37). In addition to the recapitulation of *in-vivo* ablation results, we perform *in-silico* ablations to further elucidate the obtained results. In the experiments, these were not performed due to technical challenges or other aspects, e.g., separate ablation of PVC and LUA was not possible in (34). Since in the model we can ablate any subset of neurons, we perform these, and further ablations and include these results in SM (Fig S 12;).

**Figure 6:**
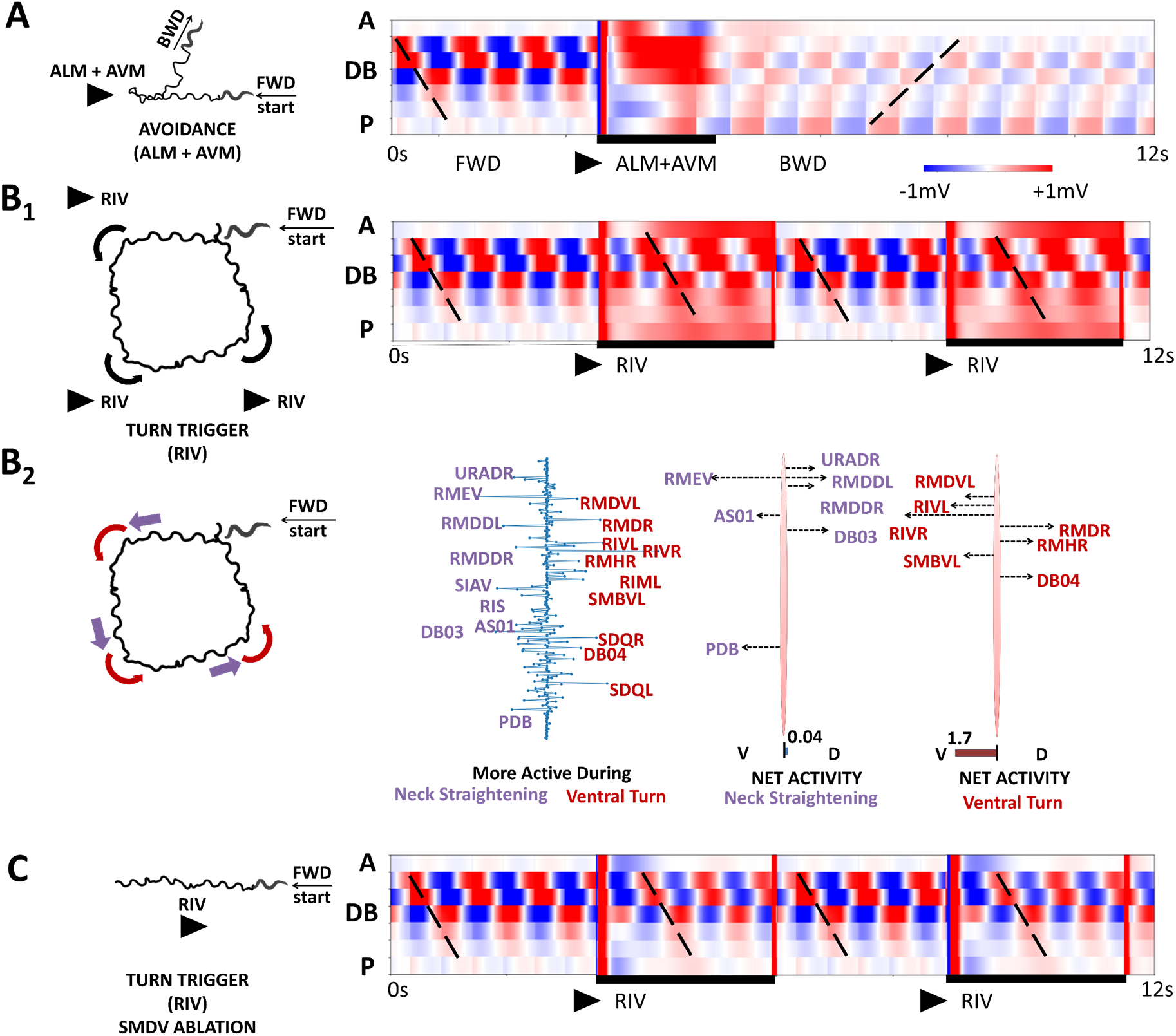
Neural impulses modify basal locomotion behavior. **A**: Avoidance behavior in which forward locomotion is interrupted by ALM+AVM impulse leading to backward locomotion (left: head trajectory, right: motor neurons (DB) activity). **B1**: RIV stimulus (2.7nA) is applied every 6 sec for duration of 3 sec during forward locomotion and rotates locomotion direction by 90°. **B2**: Dominant neurons and forces involved in rotation of locomotion direction (2^nd^ column): (i) neck straightening-purple (ii) ventral turn-red. 3rd and 4th columns: dominant motor neurons and corresponding forces they generate in ventral and dorsal directions (3rd column - neck straightening, 4th column - ventral turn). **C**: The effect of SMDV ablation: typical RIV response (rotation by 90°) is disabled.

#### 3.6.2 *In-vivo* validation of posterior gentle touch model prediction for AVA ablation

For (PLM stimulation (Gentle posterior scenario); left-bottom in Fig. 5A), *in-silico* ablations provide novel predictions: while multiple ablations are consistent with *in-vivo*, the model indicates that ablation of either (AVA) + (AVD) or (AVA) significantly alter the response such that the movement is slower than control and in about half of the simulated cases invokes *backward* movement instead of the control forward movement. Since ablation of (AVD) alone does not result in significant change in the response, the model identifies AVA interneurons to have a vital role in forward movement contradicting the classical classification of AVA as a *backward* command neuron and the reported results of (34,37). Such a finding on AVA role in forward movement has been presaged by (141) and corroborated by recent experimental works (142).

Since such response was not identified in the original experiment of gentle posterior touch response, we validate the prediction with a *novel in-vivo* experiment. We use optogenetic miniSOG method to ablate AVA neurons in ZM7198 mutants and compare their responses to gentle posterior touch with control wild type N2 animals (see Fig. 5B and SM for Videos, Methods, Behavioral Assays). Gentle posterior touch was performed mechanically with hair touching the posterior part of the body. Control *in-silico* animals exhibited sustained forward movement with average instantaneous velocity of 142 ± 22 µm/s for the duration of at least 10s after the posterior touch onset. *In-vivo* control worms exhibited matching velocities, forward postures and bearing with *in-silico* control worms.

With AVA neurons ablated, *in-vivo* worms are unable to perform sustained forward movement. Behavioral assays indicate average instantaneous velocity of 19 ± 12µm/s after the posterior touch which was in-line with the velocity of 13 ± 22µm/s reported by *in-silico* AVA ablated. In addition, 61% of tested animals (16/26) performed spontaneous backward movement for the duration of at least 1s (as can be seen in Fig. 5B bottom; red color corresponds to backward). The result is in qualitative agreement with recent *in-vivo* findings by (142) and *in-silico* ablation of AVA (see side by side comparison in Fig. 5B).

### 3.8. Timed neural stimuli and neural ablation effects

The touch response case study leads us to explore extensions of neural stimulus complexity and to examine the effect of timed external stimuli impulses while the worm is performing locomotion. Here we consider several case studies. The first case study is avoidance (Fig. 6A), which can be induced when forward locomotion is interrupted by ALM + AVM stimulus via anterior touch (143). In such scenarios, *C. elegans* reacts to the stimulus by stopping and reversing. We examine this scenario by initiating forward locomotion and after 6 seconds, applying ALM + AVM neural stimulation for 2 seconds. As we show in Fig. 6A, ALM + AVM neural stimulus indeed produces body dynamics similar to the avoidance behavior observed *in-vivo*, where the stimulus disturbs forward locomotion followed by initiating backward locomotion. Inspection of neural activity of motor neurons (DB neurons are A→P ordered in Fig. 6A) indicates that the stimulus induces a change in the directionality of the traveling wave of neural activity from A→P to P→A. The transition is marked through high constant activity in the anterior motor neurons.

An additional case study that we consider is the sharp ventral turn that occurs during reorientation in the direction of locomotion (Fig. 6B). The RIV motor neurons synapse onto ventral neck muscles and have been implicated in the execution of the ventral turn (48,128). To examine the role of RIV, we stimulate RIV neurons every 6 seconds for a duration of 3 seconds, while the worm is performing forward locomotion. As we show in Fig. 6B1, each RIV stimulus causes a sharp ventral bend of the head leading to a rotation of forward locomotion course by approximately 90° while sustaining locomotion in the forward direction. Neural activity indicates that the turn corresponds to a bias added to the membrane potential activity of oscillating motor neurons. The rotation of the body is exhibited by two posture states: (i) neck straightening followed by a (ii) ventral turn. These states are observed in experimental studies of the escape response as well (143). We investigate these states by performing SVD on neural activity in each state and identify dominant neurons associated with the activity. We then compute the force magnitude resulting from dominant motor neurons activity (Fig. 6B2). We find that during neck straightening state, dorsal and ventral forces tend to be balanced and cancel each other out, while in the ventral turn state there is a strong ventral force acting on muscle segments. Such analysis reveals neural participation on the cellular level in each state and how neural activity is superimposed to create a particular posture.

As in the touch response studies, systemic ablation can be used to further analyze the observed behaviors. Here we utilize *in-silico* combinatorial ablation of neurons to seek which neurons would be most correlated with this behavior. We select all pairs of neurons (R and L) from the group of neurons directly connected to RIV and separately ablate each pair. The analysis shows that the ablation of SMDV causes the most prominent change in dynamics, where it disables the turn behavior and causes the body to continue with forward movement, see Fig. 6C. Neural activity in the case of SMDV ablation is similar to neural activity during forward movement, suggesting that both SMDV and RIV neurons are required to facilitate a sharp turn.

### 3.9. Model variations studies

#### 3.8.1 Empirical variations for investigation of simulated behaviors

Novel experimental data shows that there are additional *higher-order* properties that play role in neural activity and behavior such as spiking neurons, novel connectomics data, and extra-synaptic connections etc (16,48,109,110,112,144). The proposed base model, through modWorm, supports the incorporation of these additional properties and potential future variations. We therefore incorporate these model variations to explore the refined fits of eigenworm characteristics for the forward and backward movements in the touch response case study.

The variations we consider are (1) neurons with non-linear ion channels, (2) variation of the connectome, and (3) tyramine-gated chloride channels (LGC-55). Each of these variations was recently proposed in experiments to have a potential role in mediating locomotion (16,48,112,144). For variation (1) a group of (AWA, AVL) neurons was shown to exhibit all-or-none spiking action potentials (105). The spikes are mediated by non-linear voltage-gated calcium and potassium channels and were initially observed in AWA sensory neurons, but recent experiments found such channels facilitate action potentials in enteric motor neurons (AVL) (112). While both AWA and AVL are not known to be directly associated with locomotion, we include these variations to demonstrate incorporations of additional biophysical dynamics to the base model and their possible roles in locomotion. For variation (2), we consider variation of the connectome to an up-to-date electron microscopy reconstruction that are based on the analysis of both new and published electron micrographs (16). The updated connectomics include additional connections for both gap and synaptic connectomes across the entire *C. elegans* nervous system that were previously missing or inaccurate in the old hermaphrodite connectome data (13). Lastly, for variation (3), LGC-55 is expressed by neurons that are post-synaptic to tyraminergic motor neurons. These neurons receive inhibitory signals from tyraminergic motor neurons and are known to inhibit head movements and forward locomotion during escape response (48). In modWorm, incorporating these variations into the base model can be done by modifying intra-cellular module to incorporate new individual neuron channels (variation (1)), extra-cellular modules (gap and synaptic currents) with updated connectivity weight matrices (variation (2)), or updated neuron polarity matrix (variation (3)) (see SM and Fig. S 14).

Figure 7 and Table 1 describe the error comparison of forward and backward locomotion eigenworm coefficients (*in-vivo* vs model) obtained with base model simulations versus the models with variations. Variation (1), selected neurons having non-linear channels appear to have slightly higher error (14.8 ± 2.1%) compared to that of the base model (13.1 ± 0.8%), where the increase in error is mostly due to AVL modeled by HH-model with non-linear channels. Variation (2), updating the connectome data to the dataset published in (16) on the other hand, significantly decreased the error between the variant model and *in-vivo* coefficients (5.6 ± 0.7%). Especially the backward movement *in-vivo* coefficients were within the confidence interval (*P = 95%*) of model obtained coefficients, indicating a closer match. From the incorporation of LGC-55, variation (3), we observe that errors for both forward and backward locomotion do not change significantly from their base counterparts (12.8 ± 0.8%). This is expected since LGC-55 was found to be primarily associated with head turning behavior during pirouette maneuver, e.g., omega turn, rather than basal locomotion (48,144). Combining all three variations decreased the error for forward (8.8 ± 2.3%) but slightly increased the error for backward locomotion (13.0 ± 0.5%). These results suggest that the effects of variations are not additive and indicate the existence of additional processes that may contribute to shaping *in-vivo* locomotion postures, and the importance of connectomes on the body dynamics.

**Figure 1:**
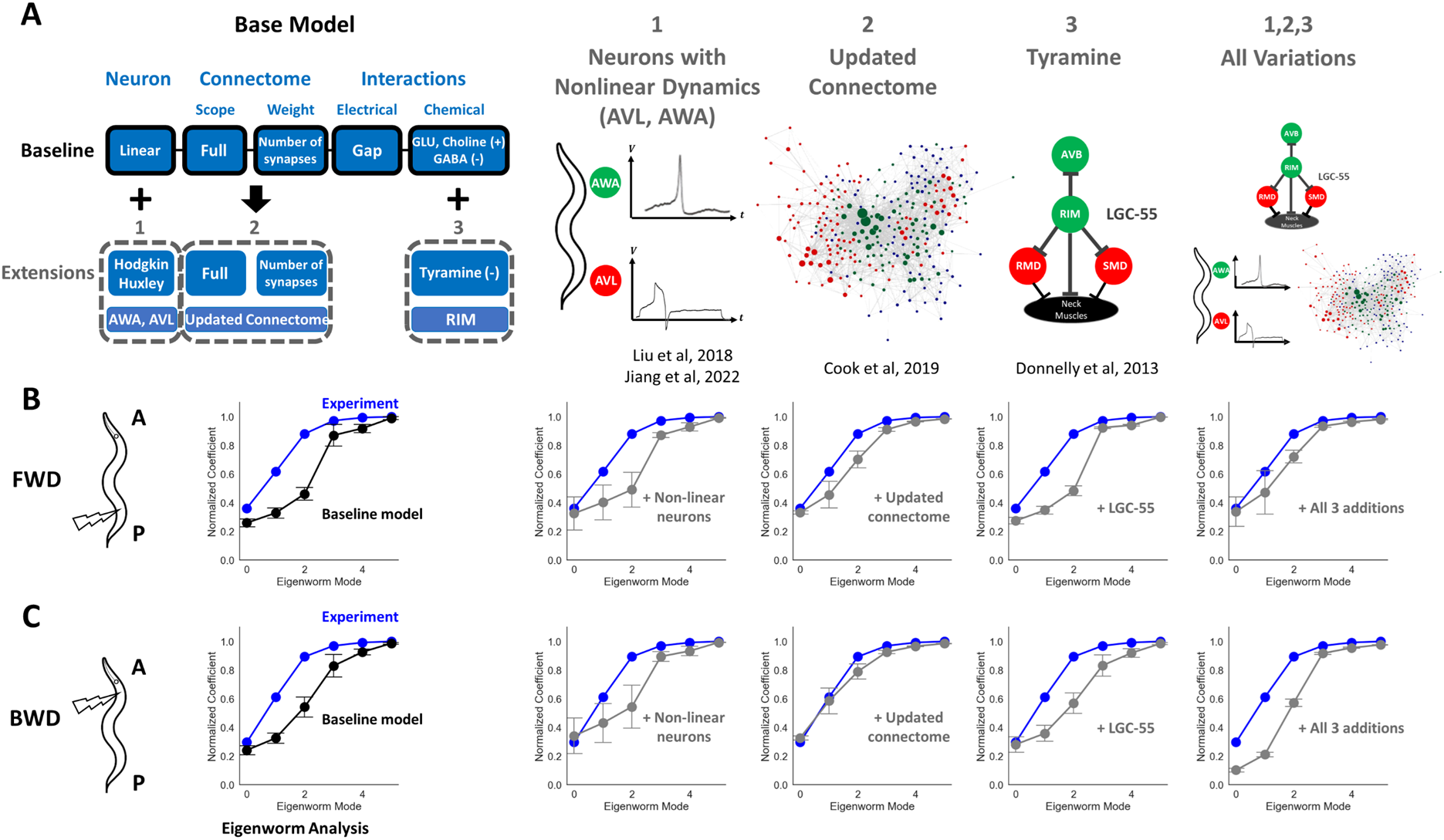
Model variations and their effects on eigenworm coefficients obtained from simulated FWD and BWD locomotion. **A**: Graphic illustrations of considered extensions to the model – from left to right: Neurons with “known” non-linear channels: AWA and AVL, updated connectome mappings, Tyramine gated chloride channels (LGC-55), and combination of all three additions. See Supplementary Materials for detailed implementations. **B**: Comparison of cumulative eigenworm coefficients during FWD locomotion between experiment (blue) vs base model (black) and each of the model extension (grey, N = 10, p = 0.05). FWD locomotion is simulated in the model by injecting a pulse of 3nA (±10% variations each trial) of current into PLML/R followed by entrainment of proprioceptive feedback. For updated connectome, the synaptic weights are randomly varied by ±10% each trial. **C**: Comparison of BWD locomotion between experiment vs base model (black) and each of the model extension (grey, N = 10, p = 0.05). BWD locomotion is simulated in the model by injecting a pulse of 6.8nA and 3nA of current into ALML/R, AVM (±10% variations each trial) respectively with proprioceptive feedback. Synaptic weights are randomly varied by ±10% each trial for updated connectome addition.

**Table 1:**
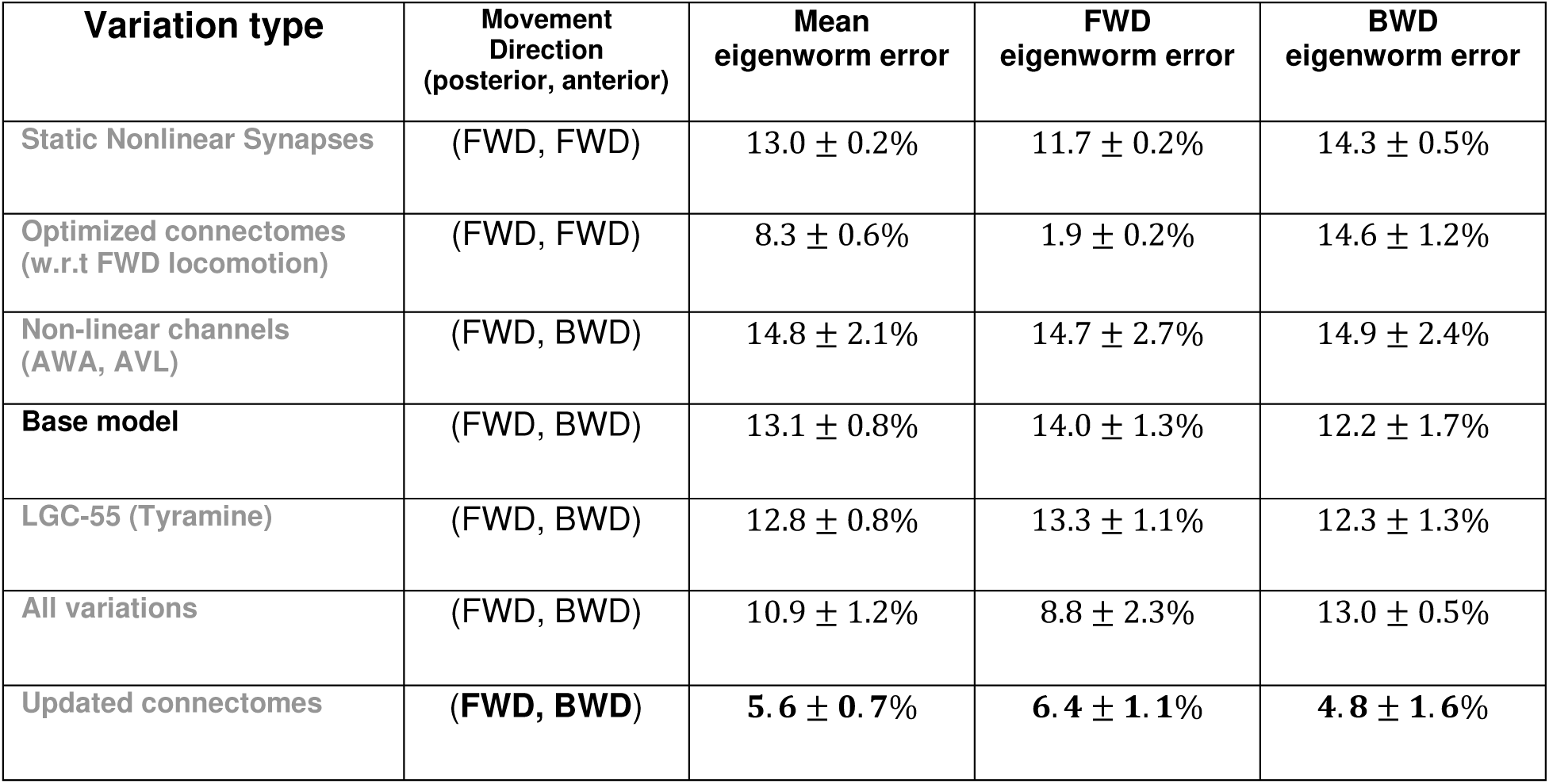
For each model variation (row), movement directions with respect to posterior/anterior touch stimuli and normalized eigenworm coefficient error vs experiment are shown. Model variations are first sorted with respect to correct movement direction associated with posterior and anterior touch (FWD, BWD) which are then sorted with respect to mean eigenworm coefficient error in descending order.

#### 3.8.2. Model optimization

Following the improvements of simulated behavior with variation (2), updated connectome data, we asked how further optimization of the connectome parameters with respect to specific behavior could affect the overall simulated behavior errors. Such task-optimized neural parameters have been successful at predicting experimental neural activity and behaviors for different organisms including Hydra, *C. elegans*, and fruit fly (73,99,115). We used a Genetic Algorithm to optimize individual synapse strengths of updated connectome data from (16) (both gap and synaptic) with respect to *in-vivo* forward locomotion eigenworm coefficients. The optimization reduced the normalized coefficient error from 5.8% to 1.9% for simulated forward locomotion (Fig. S13) (See SM for optimization procedure). To validate whether these parameters are generalizable to other types of locomotion, we apply neural stimulus associated with backward locomotion and evaluate its errors with respect to associated *in-vivo* eigenworm coefficients. This resulted in a significant increase in the error from 4.8% to 14.6% and changed the movement direction from backward to forward (Table 1, row 2). The results indicate that neural parameters (e.g., connectome mappings) optimized to specific behavior are not necessarily generalized to other behavior types or indicate biophysical parameters.

#### 3.8.3. Alternative module for neural dynamics

In addition to model optimization with respect to neural parameters, we conduct variations on model dynamics equations to study its effects on simulated behaviors. We specifically modify the synaptic dynamics term I^syn^ so that the transmitters release gating variable s(t) is calculated with a static nonlinearity (e.g., sigmoid) dependent only on membrane potential (See SM for details). Such a variation would be more comparable to artificial neuron activation function and is a simpler approach than the original synapse model where the gating variable s is governed by its own differential equation similar to Huxley-Hodgkin channel (145). The results show that while the simulated behavior under such variation has similar eigenworm coefficient errors as the base, it results in abnormality in movement properties where both posterior and anterior touch responses result in forward locomotion and the locomotion speed is significantly slower (Table 1, row 1). The results suggest that the detailed synaptic transmission model may play an important role shaping the simulated locomotion behavior closer to *in-vivo*.

## 4. Discussion

Our study provides a novel modular computational approach: *modWorm*, and subsequent neuro-mechanical model of *C. elegans* to explore the interaction between its nervous system and behavior. The proposed base model includes a total of 7 sensory-neuro-mechano-environmental modules incorporating the connectome of the full somatic nervous system, its response to stimuli and effects of neural activity on body postures and proprioception. Applications of simple stimuli in the model show that the structures of the connectome and polarities set specific movement patterns enabled by neural dynamics and biomechanics. We show that the transformations between the different constituent modules are in the form of dynamic mappings (43,45). This appears to be essential to determine whether neural activity can generate coherent body movements and cannot be resolved without incorporating the complete connectome. We introduce methods for forward integration (stimulus to muscles) and inverse integration (muscles to neural activity) allowing us to find correlation between neural stimuli and muscles and close the loop between them through proprioceptive feedback. We test the model by applying spatially travelling wave forces along the body followed by injecting constant currents into neurons in the touch response circuit. We observe that stimulation of a few neurons (sensory and inter neurons) in these circuits can generate coordinated movements consistent with direction and patterns as in *in-vivo* experiments. We then examine the effect of proprioceptive feedback and show that feedback with a time delay can entrain, smooth, and sustain locomotion initiated by neural or external force stimuli. We further test our model against previous touch response *in-vivo* experiments which used ablations to identify key neurons involved in the responses. We repeat these ablations and perform additional ablations, that were not performed in those studies or infeasible *in-vivo*, to further elucidate the roles of participating neurons in these circuits.

The model’s ability to generate robust directional locomotion allows for identifying functional neural circuits and pathways associated with timed neural stimuli during locomotion. We show examples of timed neural stimuli applications during locomotion which give rise to intricate locomotion patterns and orchestrated behaviors. Specifically, we trigger behaviors such as avoidance and sharp turns through ALM+AVM and RIV neural impulses. We demonstrate that *in-silico* ablation surveys can identify neurons participating in the sensorimotor pathway of these behaviors, e.g. SMDV neuron in RIV impulse pathways.

*In-vivo* validation indicates the potential of the model to inform and complement experimental studies. In depth analysis of motion, however, shows that the characteristics of locomotion, such as eigenworm coefficients, are similar but do not precisely match with *in-vivo* coefficients (Fig. 7 Left). This is unsurprising, since the base model, by design, includes only base processes of individual neural dynamics and connections to reflect the dominant dynamic patterns and behavior. We thus utilize modWorm and introduce systemic variations to the base model to explore which additional aspects contribute to locomotion. We incorporate variations based on experimental findings such as neurons with individual non-linear channels (AWA, AVL), updates to connectome data, and synaptic channels driven by additional neurotransmitter (e.g., tyramine), that can assist in investigating which aspects contribute to better fit in locomotion metrics between *in-vivo* and the model. The results indicate that the global network properties such as connectomes may play primary roles in modulating locomotion body postures. In addition to empirical variations, we introduce theoretical variations such as task-optimized synaptic parameters and alternative model for synaptic dynamics, to elucidate their effects on simulated behaviors. Beyond the considered variations, information of the connectome, neural dynamics and processes such as monoamines signaling and neuropeptides activity modulation are rapidly become available (16,109–112,144). Moreover, modeling methodologies of biomechanics are starting to adopt three dimensions and continuous models (95,146). Investigations of the additional effects of such components could further indicate behaviors and processes that are currently not included in the base model and identify additional dynamics observed *in-vivo* such as generative spontaneous behaviors.

Beyond model optimization considered in the study, ModWorm can be used in conjunction with additional machine learning methods such as deep learning algorithms to further inform model investigations (97). For example, modWorm could be used to generate a large amount of simulation data necessary to train machine learning methods that aim to interact with the biophysical neural or body model (105,147). Leveraging the modular structure, it would also be possible for modWorm to treat parameters of biophysical modules as *learnable* (e.g., connectome weights, neuron polarity mapping) and optimize them with respect to particular neural and body task with gradient-based deep learning training methods (e.g., back-propagation through time) (148). Indeed, such approaches have already been applied to *C. elegans* and other organisms to achieve model fitting or inference of neural functions (99,102,115,149). These optimizations, however, need to be performed along with generalization requirements. Our results with connectome parameter optimization only for a subset of behaviors suggest potential simulation inaccuracies in other behaviors. Additional optimization methods (e.g., multi-objective cost function) in conjunction with gradient-based training algorithms thus can be investigated to mitigate such discrepancies. ModWorm employs community standard Python methods for constructing models (e.g., Python class) and data format (e.g., NumPy arrays). The framework is thus expected to integrate well with existing machine learning libraries (e.g., PyTorch) which adapt similar model and data structures with additional features (e.g., auto-differentiation) necessary for advanced training algorithms (150).

In conclusion, modWorm allows simultaneous simulations and variations of the nervous system, muscles and body could assist in identifying, enumerating, and classifying sensorimotor pathways. Combination of such computational studies with empirical examination and adaptation of the model may extend understanding of currently known neuromechanical functions and potentially lead to the unravelling of novel *C. elegans* brain circuits responsible for locomotion.

## Data Availability

modWorm code and generated studies reported here, manuals and tutorials demonstrating usage of the model and its variation, are available at Github repository:

https://github.com/shlizee/modWorm (to be made public upon publication).

Along with the code, a blog describing the studied scenarios is available as part of the Github repository. The blog will be open, and the community will be invited to contribute to it.

https://shlizee.github.io/modWorm/

## Acknowledgments

The authors are thankful to the reviewers for their constructive comments. ES acknowledges the support of NSF DMS-1361145, NSF IIS-2113003 and Washington Research Fund. MA acknowledges the support of NIH RO1 GM140480. ES, JK, and JS acknowledge the support of the departments of Applied Mathematics and Electrical & Computer Engineering, the Center of Computational Neuroscience (CNC), and the eScience Center at the University of Washington in conducting this research. We thank Linh Truong and Rahul Biswas from the department of Electrical & Computer Engineering for reviewing the framework codebase.

## Notes

http://faculty.washington.edu/shlizee/dynamicworm/

### Competing Interest Statement

The authors have declared no competing interest.

### Summary of Updates

added two references on AVA role that support reported findings

