## Supplementary Materials for "Modular integration of neural connectomics, dynamics and biomechanics for identification of behavioral sensorimotor pathways in *Caenorhabditis elegans*"

Jimin Kim[1], Jeremy T. Florman[3], Julia A. Santos[2], Mark J. Alkema[3], Eli Shlizerman[1,2,*]
[1] Department of Electrical and Computer Engineering, University of Washington, Seattle, WA
[2] Department of Applied Mathematics, University of Washington, Seattle, WA
[3] Department of Neurobiology, UMass Chan Medical School, Worcester, MA
[*] Corresponding Author

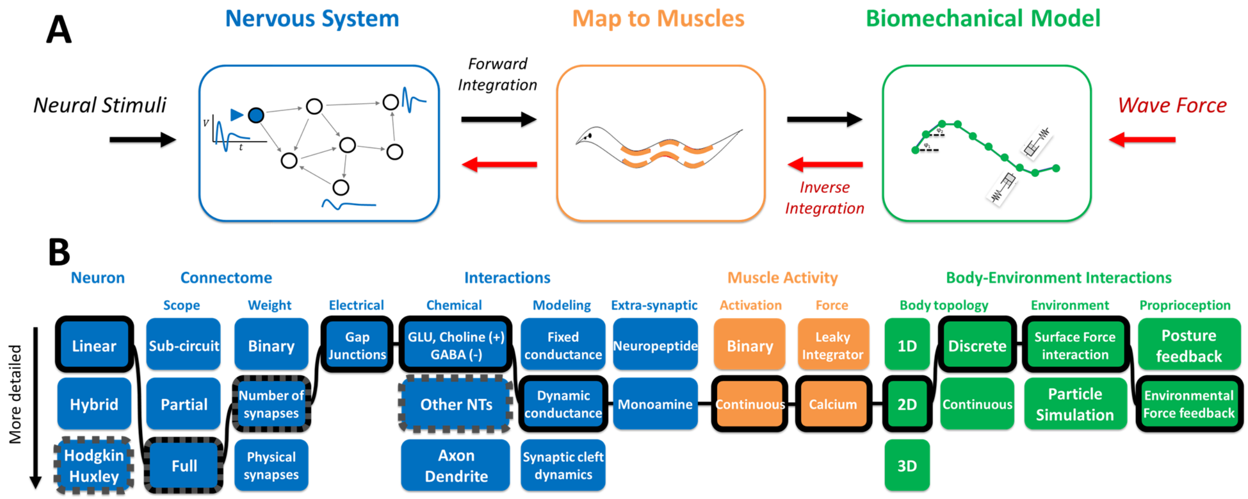


**Figure 1: Constructing *C. elegans* neuro-mechanical model**. **A:** From left to right, Modeling the nervous systems as a dynamical system encompassing the full somatic connectome including graded ion-channel and connectivity neural dynamics, mapping neural dynamics to dynamic muscle impulses and forces, mapping muscle forces to a biomechanical model that incorporates body responses and interaction with the environment. Neural stimuli are integrated forward to resolve body movements (black arrow). External forces are propagated in an inverse direction to resolve corresponding neural dynamics (red arrow). **B:** From left to right: Schematics of each model aspect as flowchart for the nervous system (blue), map to muscles (orange) and biomechanical model (green). Black highlighted boxes connected with solid lines are model aspects chosen by the proposed base model and boxes with dark gray dotted edges represent the variations considered in the paper.

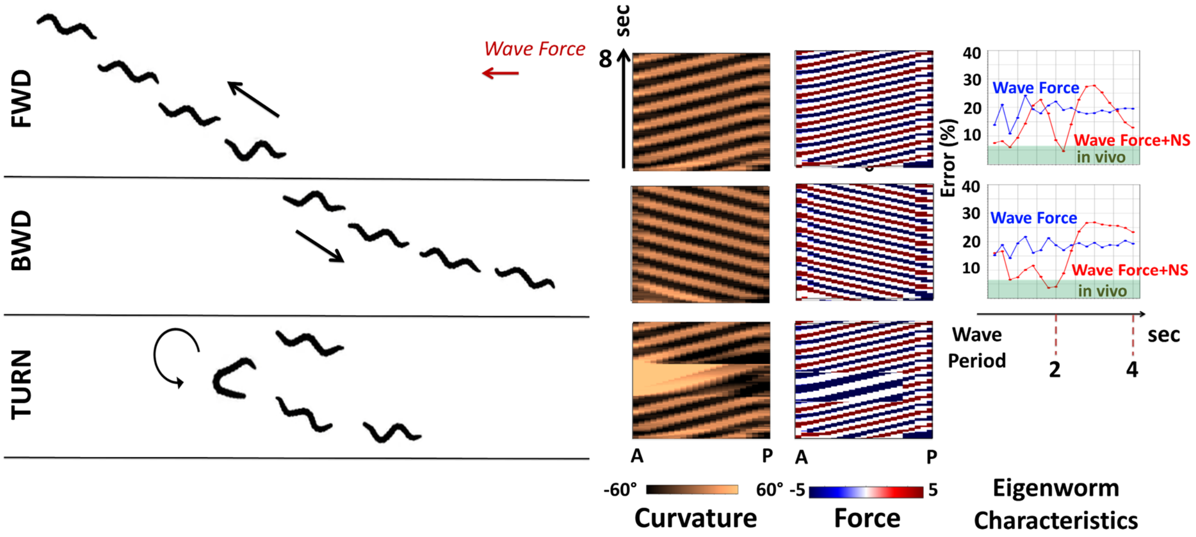


**Figure 2: Typical locomotion patterns, body curvature and force dynamics generated by three types of external wave forces, corresponding to forward, backward, 180° turn movements**. From left to right: Simulated body snapshots driven by external wave force with minimal eigenworm posture error where each snapshot is sampled every 2 seconds for forward, backward and turn dynamics (Also see SM videos), associated body curvature and muscle force dynamics ordered from anterior to posterior direction for 8 seconds simulation duration, eigenworm posture errors (w.r.t normalized posture coefficients) in the function of varying external wave force periods. Posture errors resulting from direct external wave force on the body are labeled as blue curve with 'Wave Force'. External wave force inverse integrated to resolve neural dynamics which are integrated forward to simulate body dynamics are labeled as red curve with 'Wave Force+NS'. 'Wave Force' curve results with ~20% mean error for both directions of the force and does not show preference to wave period. 'Wave Force+NS' appears to be selective to period and achieves minimal error for period ~ 2s of 4.6% (fwd) and 3.6% (bwd) within the CI of *in-vivo* worms (green band; 6.7%, P=0.01) for both directions of movement.

The implementation of 2D body model allows us to simulate the complete somatic nervous system simulation (279 neurons) for such excitation. See SM for more detailed descriptions of Muscles and Biomechanics Models.

**3.2. Locomotion and corresponding neural dynamics induced by external force.** In addition to integrating the model in a feed forward manner, we develop an inverse integration approach. Inverse integration computes the neural dynamics that would result from external forces applied to the body. The approach transforms external forces acting on the body to muscle activity and then inverts the activity to membrane potential. These potentials are then integrated forward to resolve the body posture (see SM for more details). We use this approach to validate our model for three basic locomotion patterns: forward, backward and turn movements. For each pattern we apply a sinusoidal force wave travelling along the body with variable frequency to infer neural dynamics associated with it. These neural dynamics are then forward integrated by the nervous system to generate the body posture. We then simulate the integration

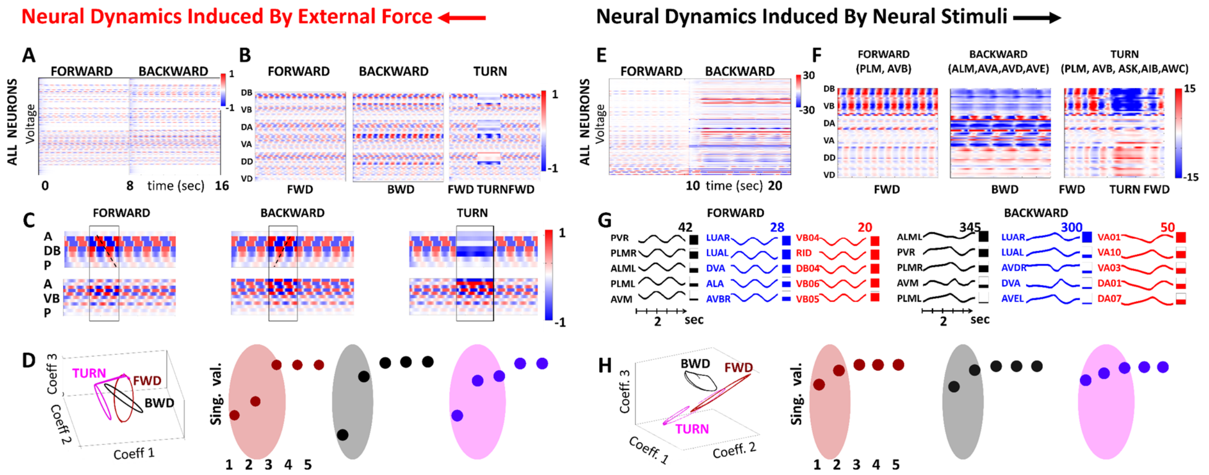


**Figure 3: Neural responses of *C. elegans* somatic nervous system to external wave forces and neural constant stimuli. A**: Color raster plot of membrane potential (difference from equilibrium) of 279 neurons inferred by inverse integration of wave forces to corresponding neural dynamics. Neural responses generated for 16 sec: 0-8 sec: spatial wave force generating forward movement; 8-9: transition; 9-16 sec: spatial wave force generating backward movement. **B**: Color raster plots of membrane potential of motor neurons for forward, backward and turn wave force profiles. **C:** Color raster plots of membrane potential of Ventral and Dorsal type B motor neurons for forward, backward and turn wave force profiles **D**: Evolution of temporal coefficients during forward, backward and turn neural responses (red, black, magenta). Temporal coefficients are associated with PC modes from SVD analysis of all three responses (i.e. projected to a common space of PC1-PC3). **E:** Color raster plot of membrane potential of optimal forward and backward current; 0-10s: forward (0.7nA into PLMR/L, 1.3nA into AVBR/L) 10-15s: transition; 15-25s: backward (2.8nA, 1nA, 0.5nA, 0.5nA into ALMR/L, AVAR/L, AVDR/L and AVER/L respectively). **F:** Color raster plots of membrane potential of motor neurons for forward, backward and turn stimulations (compare with Figure 3B). **G:** Top 5 neurons (which have largest elements in PC1 mode) from each group (sensory, inter and motor) of neurons for forward (left) and backward (right) stimuli. **H**: Evolution of temporal coefficients during forward, backward and turn neural responses (red, black, magenta); compare with Figure 3D.

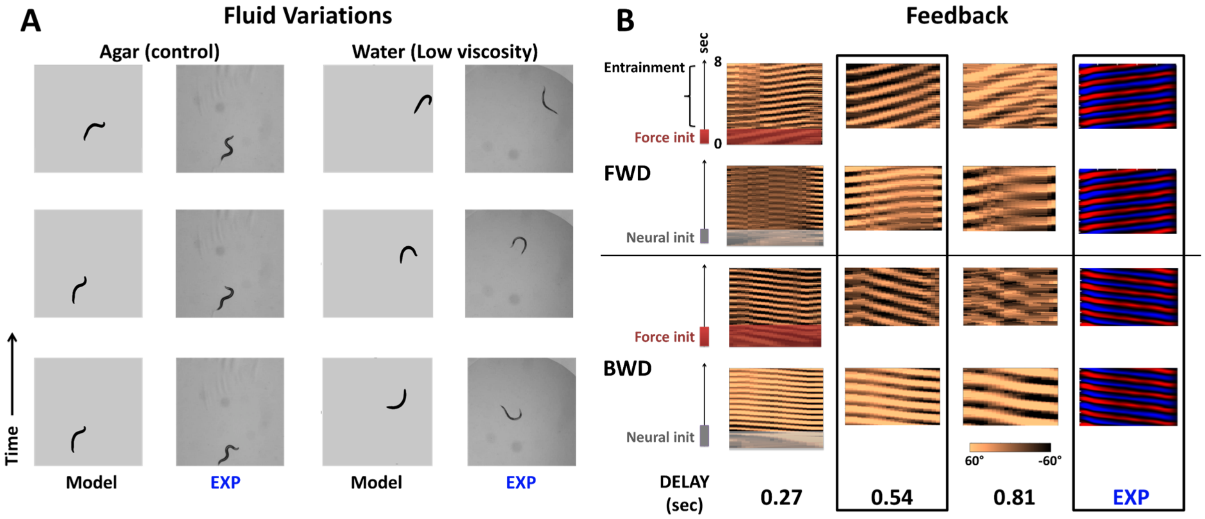


**Figure 4: Appropriate fluid parameters and proprioceptive feedback and facilitate sustained locomotion. A**: Surrounding fluid of model and experiment (134) are varied between agar (high viscosity, columns 1,2) and water (low viscosity, columns 3,4) respectively, to study their effects on locomotion. In the model, viscosity and fluid density are reduced from 10mPa/s, 1g/cm$^3$ to 1mPa/s, 0.7g/cm$^3$to emulate water droplet in experiment. For each column, 3 locomotion snapshots are taken at 1 second apart. **B**: Feedback is initiated by a wave force (rows 1,3) or neural stimuli (rows 2,4) (see also SM Videos). Columns 1-3 display various feedback time delays, modeling the environmental reaction time of producing external forces on the body, and the curvature profiles they produce and compared to column 4, experimentally recorded curvatures (adapted from (125)). Time delay of approximately 0.5 sec produces optimal forward and backward locomotion which is close to experimental locomotion (highlighted by dashed border).

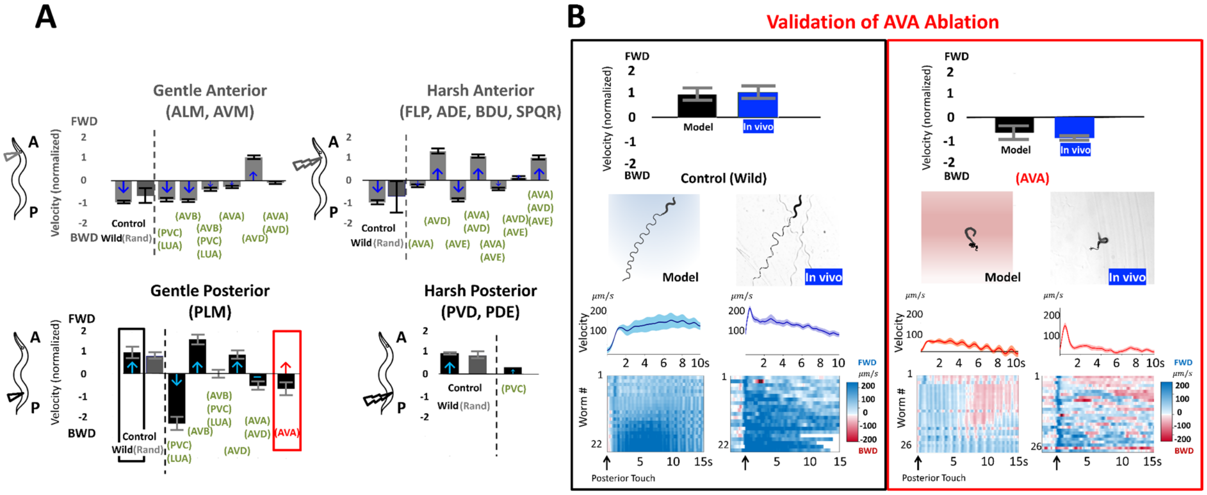

**Figure 5: Validation, recapitulation, and prediction of locomotion behaviors for touch responses. A**: Body velocities for neural stimulation associated with Top: Anterior Gentle (left) and Harsh (right) touch responses Bottom: Posterior Gentle (left) and Harsh (right) touch responses. Velocity amplitude is normalized according to the Control Wild (black label) locomotion and direction up/down are chosen as FWD and BWD direction respectively. Recapitulated velocities following ablation experiments in (34, 37) (green label) are compared to the Control Wild velocities. Arrows indicate experimental *in-vivo* observations and color indicates match of model with published experimental observations (blue: match; red: mismatch) **B**: *In-vivo* validation of Gentle Posterior Touch model prediction for ablation of AVA. Model predictions are compared with *in-vivo* assays for control (left) and AVA ablated (right). Model and *in-vivo* responses at and after stimulus onset are compared with characteristics of (top to bottom) normalized velocities, response locomotion paths, instantaneous velocities after onset, individual instantaneous velocities (each row is a worm) color plot (blue; positive velocity (fwd); red: negative velocity (bwd)).

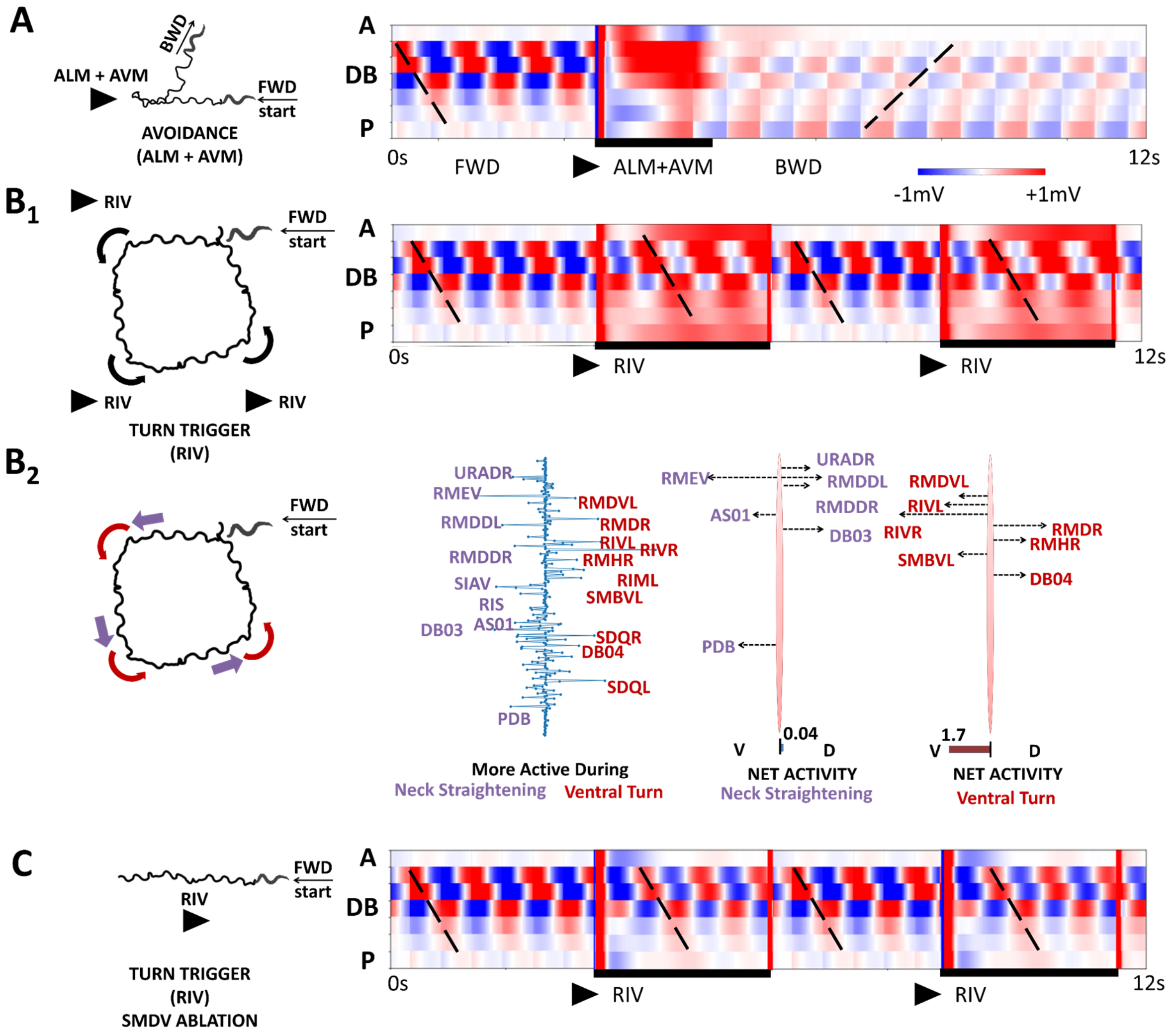


**Figure 6: Neural impulses modify basal locomotion behavior. A**: Avoidance behavior in which forward locomotion is interrupted by ALM+AVM impulse leading to backward locomotion (left: head trajectory, right: motor neurons (DB) activity). **B1**: RIV stimulus (2.7nA) is applied every 6 sec for duration of 3 sec during forward locomotion and rotates locomotion direction by 90°. **B2**: Dominant neurons and forces involved in rotation of locomotion direction (2$^{nd}$ column): (i) neck straightening-purple (ii) ventral turn-red. 3rd and 4th columns: dominant motor neurons and corresponding forces they generate in ventral and dorsal directions (3rd column - neck straightening, 4th column - ventral turn). **C**: The effect of SMDV ablation: typical RIV response (rotation by 90°) is disabled.

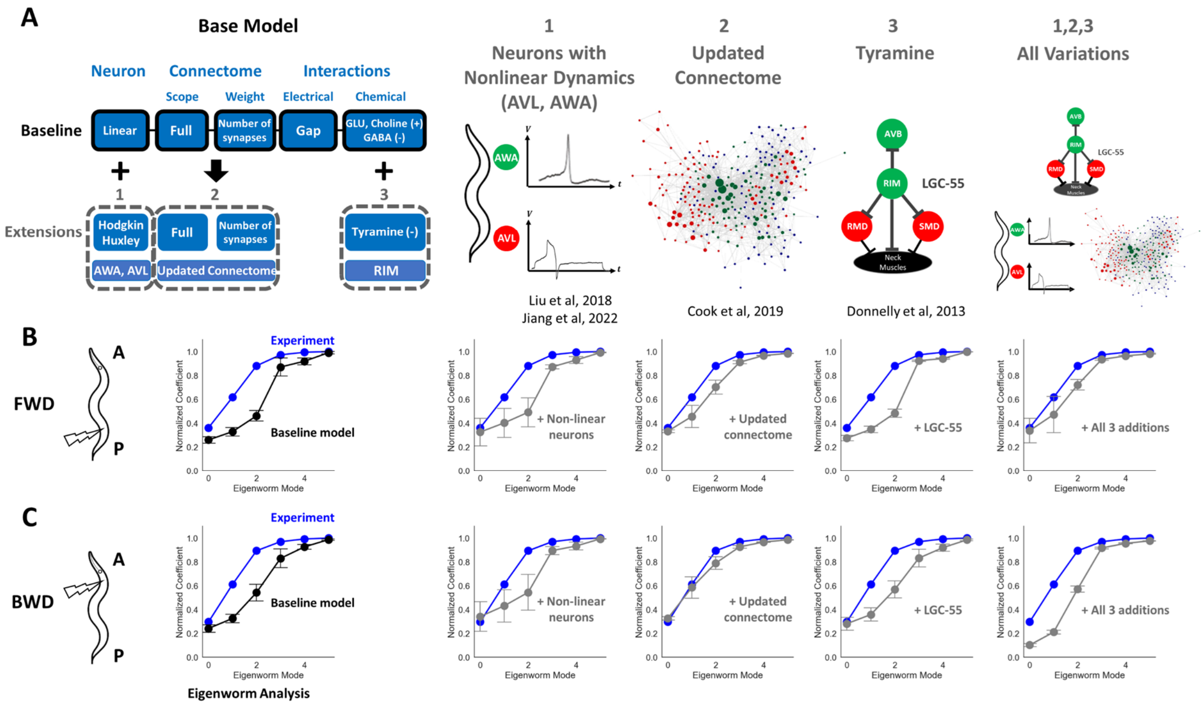


**Figure 7: Model variations and their effects on eigenworm coefficients obtained from simulated FWD and BWD locomotion. A**: Graphic illustrations of considered extensions to the model – from left to right: Neurons with “known” non-linear channels: AWA and AVL, updated connectome mappings, Tyramine gated chloride channels (LGC-55), and combination of all three additions. See Supplementary Materials for detailed implementations. **B**: Comparison of cumulative eigenworm coefficients during FWD locomotion between experiment (blue) vs base model (black) and each of the model extension (grey, N = 10, p = 0.05). FWD locomotion is simulated in the model by injecting a pulse of 3nA ($\pm 10\%$ variations each trial) of current into PLML/R followed by entrainment of proprioceptive feedback. For updated connectome, the synaptic weights are randomly varied by $\pm 10\%$ each trial. **C**: Comparison of BWD locomotion between experiment vs base model (black) and each of the model extension (grey, N = 10, p = 0.05). BWD locomotion is simulated in the model by injecting a pulse of 6.8nA and 3nA of current into ALML/R, AVM ($\pm 10\%$ variations each trial) respectively with proprioceptive feedback. Synaptic weights are randomly varied by $\pm 10\%$ each trial for updated connectome addition.

(AWA, AVL) neurons was shown to exhibit all-or-none spiking action potentials (105). The spikes are mediated by non-linear voltage-gated calcium and potassium channels and were initially observed in AWA sensory neurons, but recent experiments found such channels facilitate action potentials in enteric motor neurons (AVL) (112). While both AWA and AVL are not known to be directly associated with locomotion, we include these variations to demonstrate incorporations of additional biophysical dynamics to the base model and their possible roles in locomotion. For variation (2), we consider variation of the connectome to an up-to-date electron microscopy reconstruction that are based on the analysis of both new and published electron micrographs (16). The updated connectomics include additional connections for both gap and synaptic connectomes across the entire *C. elegans* nervous system that were previously missing or inaccurate in the old hermaphrodite connectome data (13). Lastly, for variation (3), LGC-55 is

| Variation type | Movement Direction (posterior, anterior) | Mean eigenworm error | FWD eigenworm error | BWD eigenworm error |
|---|---|---|---|---|
| Static Nonlinear Synapses | (FWD, FWD) | $13.0 \pm 0.2\%$ | $11.7 \pm 0.2\%$ | $14.3 \pm 0.5\%$ |
| Optimized connectomes (w.r.t FWD locomotion) | (FWD, FWD) | $8.3 \pm 0.6\%$ | $1.9 \pm 0.2\%$ | $14.6 \pm 1.2\%$ |
| Non-linear channels (AWA, AVL) | (FWD, BWD) | $14.8 \pm 2.1\%$ | $14.7 \pm 2.7\%$ | $14.9 \pm 2.4\%$ |
| **Base model** | (FWD, BWD) | $13.1 \pm 0.8\%$ | $14.0 \pm 1.3\%$ | $12.2 \pm 1.7\%$ |
| LGC-55 (Tyramine) | (FWD, BWD) | $12.8 \pm 0.8\%$ | $13.3 \pm 1.1\%$ | $12.3 \pm 1.3\%$ |
| All variations | (FWD, BWD) | $10.9 \pm 1.2\%$ | $8.8 \pm 2.3\%$ | $13.0 \pm 0.5\%$ |
| Updated connectomes | (**FWD, BWD**) | $\mathbf{5.6 \pm 0.7}\%$ | $\mathbf{6.4 \pm 1.1}\%$ | $\mathbf{4.8 \pm 1.6}\%$ |

**Table 1:** For each model variation (row), movement directions with respect to posterior/anterior touch stimuli and normalized eigenworm coefficient error vs experiment are shown. Model variations are first sorted with respect to correct movement direction associated with posterior and anterior touch (FWD, BWD) which are then sorted with respect to mean eigenworm coefficient error in descending order.

### SUPPLEMENTARY MATERIAL

Modular integration of neural connectomics, dynamics and biomechanics for identification of behavioral sensorimotor pathways in *Caenorhabditis elegans*

Jimin Kim[1], Jeremy T. Florman[3], Julia A. Santos[2], Mark J. Alkema[3], Eli Shlizerman[1,2,*]
[1] Department of Electrical and Computer Engineering, University of Washington, Seattle, WA
[2] Department of Applied Mathematics, University of Washington, Seattle, WA
[3] Department of Neurobiology, UMass Chan Medical School, Worcester, MA
[*] Corresponding Author

#### *I.* MODULAR INTEGRATION MODELING FRAMEWORK

We develop the modeling framework: *modWorm*, which we utilize to construct the base neuro-mechanical model of *C. elegans.* The framework is developed in Python and Julia programming languages and integrated with libraries such as NumPy, SciPy, DifferentialEquations.jl, and Jupyter Notebook for high performance simulations and intuitive interface (1–6). modWorm defines three model classes according to their scopes and scales: i) Module, ii) System, iii) Model. These three classes are inclusive to one another such that Module ⊂ System ⊂ Model where a Module describes individual biophysical process within a system (e.g., individual ion channel current, synaptic current), System describes a specific system within the organism (e.g., nervous system, muscles, body) and Model describes the full system in its entirety described by the integration of Systems (e.g., neuro-mechanics). Such modular modeling framework has also been adopted by popular modeling tools for nervous systems such as NeuroML (7,8).

All three model classes have two main components: parameters and dynamics equation space, which are fully customizable and describe the input-output relationship of the model. As the smallest model class, Modules serve as building blocks for larger model classes. Each Module's parameter and dynamics equation spaces are either pre-defined by the framework or created by the user. All pre-defined Modules' parameter spaces are fully configurable. A System is constructed by simply calling and combining the existing Modules to build its parameter and dynamics equation spaces (Fig. S1). Different Systems can be further combined into a Model where users can configure how inputs and outputs of Systems interact during simulations. The Model and its simulated dynamics can be further analyzed and visualized using the framework provided plotting and animation tools. The framework also allows the constructed model to be saved as a Python class object for future reusability.

#### *II.* *C. ELEGANS* NEURONAL MODEL

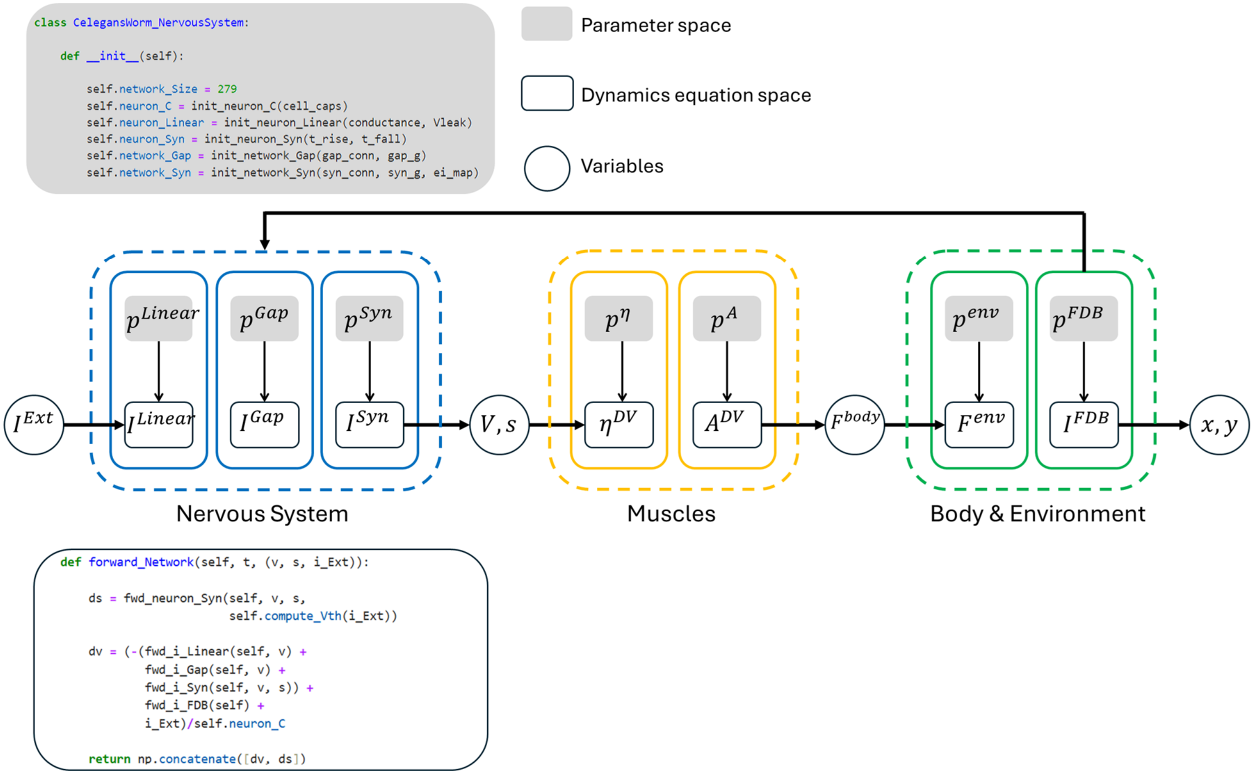


**Figure S1: Base *C. elegans* neuro-mechanical model implemented in modWorm.** The model is comprised of individual modules (solid line boxes) each with parameter and dynamics equation space. Modules are combined to form systems (dotted line boxes) which are then integrated to form a model. For the nervous system, Python code snippets outlining the example definitions of parameter and dynamics functions are shown. From left to right, neural stimuli $I^{Ext}$ is fed to the nervous system consisting of 3 modules (linear leak channel, gap, synaptic current) which outputs voltage $V$ and synaptic activity variable $s$. $V$ and $s$ are then used as inputs to muscles system to calculate muscle calcium activities $\eta^{DV}$ across dorsal-ventral muscles which are then translated to muscle activations $A^{DV}$ and outputs muscle forces $F^{body}$. $F^{body}$ is then fed to Body & Environment system to calculate corresponding environmental force $F^{env}$and proprioceptive feedback current $I^{FDB}$ and output positional coordinates $(x, y)$ for each body segment. $I^{FDB}$ is used as a feedback input to the nervous system.

To simulate neural responses, we implement the dynamical model of *C. elegans* nervous system, which consists of structural connectivity map (connectome) and equations modeling the biophysical processes of neural responses and interactions between neurons. The connectome is defined as a graph with nodes representing neurons and edges (weighted directed) representing the number of connections and their type (synaptic or gap), for more details see (9). On top of the connectome we implement dynamical equations consisting of 3 modules representing i) individual neural dynamics and two types of interactions – ii) gap and iii) synaptic (*cholinergic, glutamerigic* and GABA), as introduced in (10). In the model each neuron response is modeled by single-compartment membrane Eqn. (1) where graded potentials (leak) and gap are modeled as functions of membrane voltage $V$ and synaptic interactions between neurons are modeled with synaptic activity variable $s$ alongside with voltage $V$ as follows:

$$C\frac{dV}{dt}(t,V) = -G^C(V_i - E_{cell}) - I_i^{Gap}(\vec{\boldsymbol{V}}) - I_i^{Syn}(\vec{\boldsymbol{V}}) + I_i^{Ext} \tag{1}$$

where

$$I_i^{Gap} = \sum_j G_{ij}^g(V_i - V_j) \tag{2}$$

and

$$I_i^{Syn} = \sum_j G_{ij}^s s_j(V_i - E_j) \tag{3}$$

The parameters in the equations are total cell capacitance $C$, leakage conductance $G^C$, leakage potential $E_{cell}$, external input $I^{ext}$. Neural interactions are represented as $I^{Gap}$ and $I^{Syn}$, defined in Eqn. (2,3) and correspond to gap and synaptic junction currents respectively. The coefficients in Eqn. (2) and (3) are defined by the connectome: $G_{ij}^g$ is the total conductivity of gap junctions between neurons $i$ and $j$, and in Eq. (3), $G_{ij}^s$ is the maximum total conductivity of the synapses between neurons $i$ and $j$ modulated by the synaptic activity variable with dynamic equation:

$$\frac{ds_i}{dt} = a_r\phi(V_i;\beta,V_{th})(1-s_i) - a_d s_i \tag{4}$$

In Eqn. (4), $a_r$ and $a_d$ are the activity's rise and decay time scale coefficients respectively, and $\phi$ is the sigmoid function:

$$\phi(V_i;\beta,V_{th}) = \frac{1}{1+\exp\left(-\beta(V_i - V_{th})\right)} \tag{5}$$

Where $V_{th}$ is the network equilibrium voltage in the function of $I_i^{Ext}(t)$ that satisfies $dV(t)/\,dt = 0$. $Vth$ is thus solved at every time step to reflect time dependent $I_i^{Ext}(t)$. For more information about the derivation of the neuronal model, biophysical processes that the equations represent and units of the terms, see (10,11) and references therein.

The set of dynamical equations representing the model is 558 dimensions (2 ODEs * 279 neurons). Furthermore, these equations are stiff and thereby solved using implicit backward differentiation formulas (BDF). Specifically, we utilize VODE and CVODE solvers provided by SciPy and DifferentiEquations.jl for Python and Julia respectively when solving these equations (12,13). The system robustness also has been tested by using higher order and stochastic methods in which small external noise has been added to the parameters and verifying that the system solutions are indeed robust to these changes (10).

For efficient simulations, integration time is optimized and approximately corresponds to **actual time of the network dynamics or less**, i.e., 1 sec of integrated neural dynamics > 1 sec of actual neural dynamics. The simulation step is by default 0.01s and can be modified by the user to an arbitrary value. The integration speed can be further improved by incorporating analytic Jacobian or running the simulation in Julia as supported by modWorm. The simulations can also be run in parallel by using the ensemble simulation provided by modWorm.

#### *III.* EXTERNAL STIMULATION

The term $I_i^{Ext}$(t) represents the external time-dependent current injected into each neuron. By replacing this term with different functions, we emulate stimulation of the nervous system. Fundamental stimulation is injection of constant current $I_i^{Ext} = stim_i$ into a subset of *i* neurons. Such input is generalized to step function input, where it is turned on for some time and then turned off. To avoid abrupt changes of values and hence discontinuities in the model equations, we implement smooth transition between on and off state using a sigmoid function, from one fixed value of the input to another one. The outcome is a smooth step function with typical full transition time of 2 sec. In addition, we use $I_i^{Ext}$(t) for time-dependent stimulations, such as periodic functions with various phases, periods, and amplitudes (smoothing is not required for these functions). Here, we generated the neural dynamics related to the touch circuit by using combinations of step functions as well as periodic stimuli (summarized in Table S1). Membrane voltages that are displayed and used for transformation to muscle activity (plotted in Figure 3) are voltage displacements between each neuron's membrane voltage value at time $t$, $V_i(t)$, and its resting state $V_{th}(t)$, i.e. $\bar{V}_i = V_i(t) - V_{th}(t)$.

| Stimulation Type | Neurons | Range (pA) | Behavior |
|---|---|---|---|
| Step | PLMR/L | 0-1500 | - |
| Step | PLMR/L, AVBR/L | 0-1500, 0-2500 | FWD |
| Step | ALMR/L | 0-5000 | - |
| Step | ALMR/L, AVAR/L | 0-5000, 0-2000 | - |
| Step | ALMR/L, AVDR/L | 0-5000, 0-2000 | - |
| Step | ALMR/L, AVER/L | 0-5000, 0-2000 | - |
| Step | ALMR/L, AVAR/L,AVER/L, AVDR/L | 0-5000, 0-1000, 0-1000, 0-1000 | BWD |

| | | | |
|---|---|---|---|
| Step | PLMR/L, AVBR/L, ASKR/L, AWCR/L, AIBR/L | 0-1500, 0-2000, 0-2000, 0-1000, 0-1000 | TURN |
| Periodic $1 + a\sin(p2\pi/t)$ | PLMR/L | a: 0-3000<br>p: 0.5-4 | FWD, BWD |
| Periodic $1 + a\sin(p2\pi/t)$ | ALMR/L | a: (0-3000) * 2.3<br>p: 0.5-4 | FWD, BWD |
| Periodic $1 + a\sin(p2\pi/t)$ | PLMR/L, ALMR/L | a: (0-3000)<br>p: 0.5-4 | FWD, BWD |

**Table S1:** Summary of step and periodic stimuli (neurons, ranges of amplitudes and parameters) applied to neurons for investigation of the touch response.

#### *IV.* STIMULI OPTIMIZATION

To determine whether step stimulations can induce coordinated movements we employ both gradient based approach (line search) and direct optimization (survey of a range of parameters) for stimulus amplitude. The gradient method is used to identify which neurons and range of stimulations can produce locomotion in a particular direction. Direct optimization is then used to accurately map these ranges. For estimation of the locomotion coordination and direction we use a simple measure of average distance passed by a segment on the body model (tail or head). We find the optimization function to be non-convex with multiple minima, which makes it difficult to find global minimum and requires multiple trials by optimization procedure to find sufficient minima using the line search.

We set the algorithm to search for step currents that optimize the distance of locomotion into a particular direction and the total squared current amplitude $\|\vec{a}\| = \sqrt{\sum_i a_i^2}$. We focus on touch circuit and examine stimuli into neurons (sensory, inter neurons) that were found experimentally to be involved in this circuit (14,15). For forward locomotion, line search shows that the space could be spanned by two currents: PLML/R and AVBR/L (with identical current into both left and right neurons). We compute the space of currents in the range of (0, 1.5nA) for PLMR/L and in the range of (0, 2.5nA) for AVBR/L, where 1nA = 1000pA. We find the minimum with lowest amplitude at PLMR/L = 0.7nA and AVBR/L = 1.3nA (total squared amplitude is $0.73nA^2$), see Fig. S2 Left. For backward locomotion, line search identifies four currents that play a critical role: ALMR/L, AVAR/L, AVDR/L, AVER/L. We fix 2 two-dimensional spaces and first find optimal parameters in AVDR/L, AVER/L space (Figure S2 Right). This resulted in AVDR/L = 0.5nA, AVER/L = 0.5nA according to our minimum with lowest amplitude. We then follow

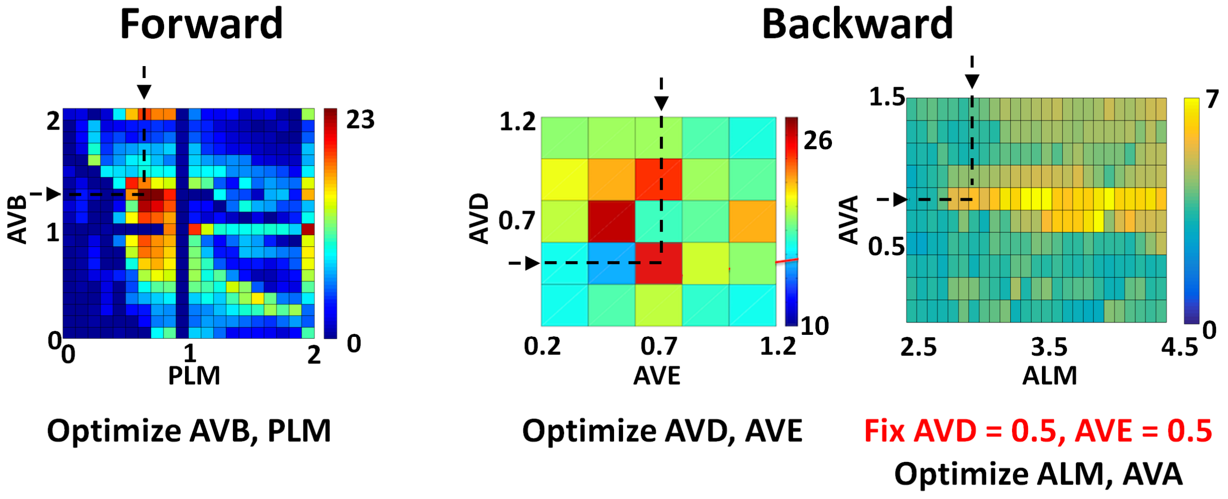


**Figure S2:** Optimization for forward and backward locomotion. Left: Optimization over two-dimensional parameter space (AVB and PLM) which results in the maximum forward distance traveled. Right: Optimization over four-dimensional parameter space (ALM, AVA, AVD, AVE) which results in the maximum backward distance. The parameter space is split into two spaces (AVD, AVE) first and then (ALM, AVA). Color corresponds to distance passed during locomotion. 1000pA = 1 is used for stimulus amplitude unit.

with a search for a minimum in ALMR/L, AVAR/L space. We find the minimum with lowest amplitude to be at ALMR/L = 2.9nA and AVAR/L = 1nA (total squared amplitude is $0.78nA^2$), see Figure S2 Right.

For turn dynamics we add the currents of AIBR/L, ASKR/L, AWCR/L neurons to forward optimal stimulus and employ a direct search to find the lowest currents that will modify typical forward neural and behavioral responses. Multiple current settings change responses in various ways. We find the lowest and most significant change when we set ASKR/L = 0.3nA, AWCR/L = 0.6nA, AIBR/L = 0.5nA and use it in Fig. 3.

#### *V.* DIMENSION REDUCTION OF NEURONAL DYNAMICS

We investigate the patterns of the *C. elegans* neural network activity during forward, backward locomotion and turn by performing singular value decomposition (SVD) on neuronal activity data (voltage difference from equilibrium) (see Fig. 3 and Table S1). We focus on analyzing ten-second simulations which were sufficient to describe the representative dynamics associated with the stimuli. In all cases, the first second of simulation data was removed to ignore the effects of the transitional forced perturbation. The SVD is applied on simulation data, which is a matrix *D* of dimensions $n \times t$ (each row corresponds to $i$th neuron, and column corresponds to a time sample $t$) and its elements are $\bar{V}_{i,t}$ (16,17). The raster plots for matrices *D* for different types of neuron groups (complete connectome, dorsal motor

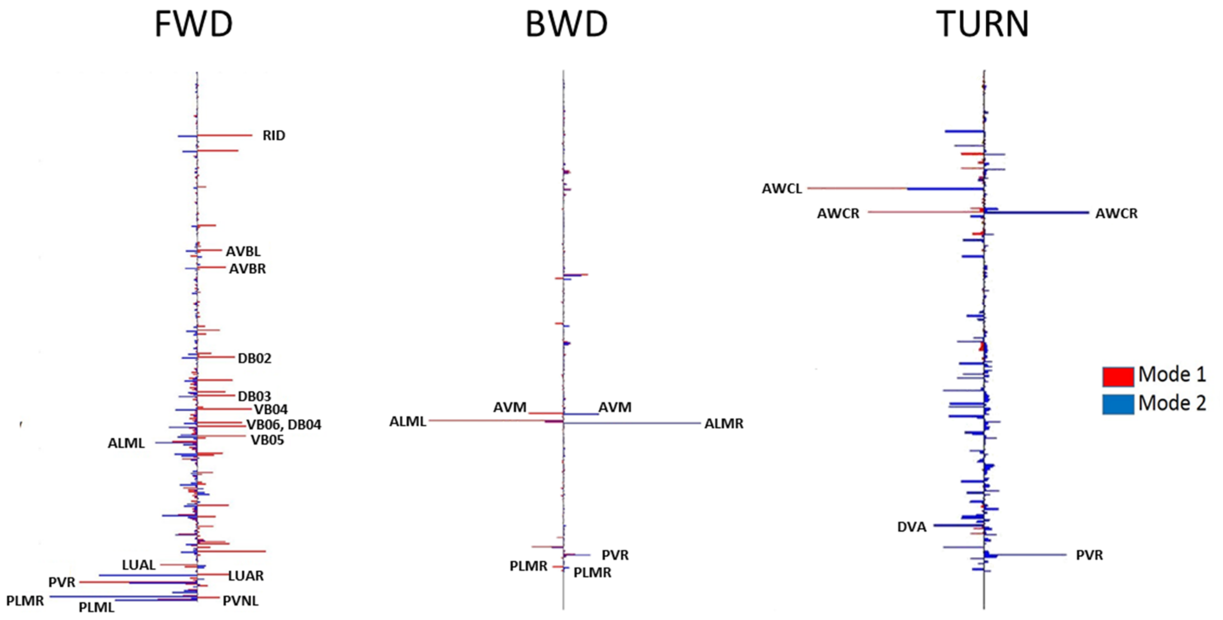


**Figure S3:** Pattern values for the first two SVD modes for neurons along the body ordered from Anterior (top) to Posterior (bottom) during forward, backward, and turning movements. Neurons in the posterior region dominate during forward motion whereas backward motion appears to be dictated by a few key neurons and turning engages anterior neurons more heavily.

neurons, ventral motor neurons) and different external inputs (FWD, BWD, TURN) are shown as color plots in Fig. 3.

The SVD decomposes $D$ into $D = \sum_{l=1}^{N} u_l \sigma_l v_l^T$, where $u_l$ are the eigenvectors of $DD^T$, representing the pattern of each mode (PC modes), $v_l$ are the eigenvectors of $D^TD$, representing the time-dependent coefficients of each mode, and $\sigma_l$ are the eigenvalues of both $DD^T$ and $D^TD$, which are the singular values that act as stretching factors. We also consider the k-mode truncated decomposition of $D$ as $D_k = \sum_{i=1}^{k} u_i \sigma_i v_i^T$ in which we take the dominant $k$ modes. For stimulation that corresponds to FWD and BWD we find that only few principal component (PC) modes are significant for the decomposition: the first PC mode explains approximately 65.24% of the energy and the first two PC modes explain about 90.94% of the energy, where the energy explained by mode k is defined as $\sum_{i=1}^{k} \sigma_i^2 / \sum \sigma^2$. Since the system is dominated by these first three PC modes, we focus on a 3-mode truncated decomposition and investigate the modes and time-dependent coefficients in this decomposition. In Fig. 3, we show the projection onto the first three modes, which corresponds to time-dependent coefficients. We also plot the absolute values assigned to each neuron in the first two modes ordered along the body of the worm in Fig, S3. We then pick the five top neurons from sensory, inter and motor neurons and plot their membranes voltages in Fig. 3 and label them in Fig. S3.

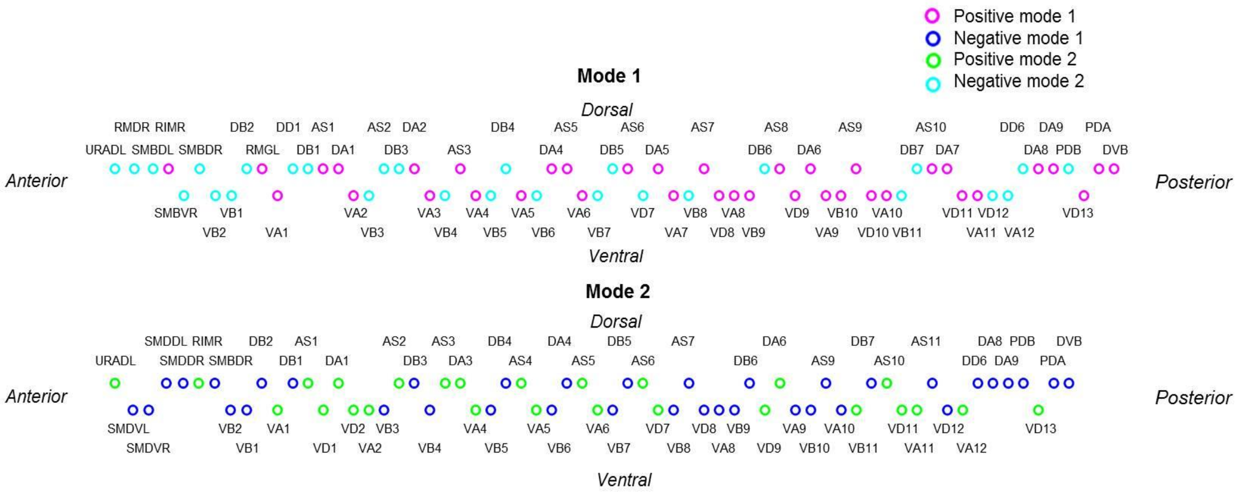


**Figure S4**: Visualization of the location and pattern values of active *C. elegans* motor neurons in the first two modes. Neurons are positioned according to their location on the AP-axis and the muscle group(s) they stimulate (dorsal or ventral). Colors indicate whether the neuron has a positive or negative pattern value in that particular mode.

In reviewing the first two modes that correspond to FWD movement in Fig. S3, it is evident that the neurons with stronger patterns values fall toward the posterior region of the body, indicating that the posterior region contains neurons that have a more significant role in *C. elegans'* forward movement. The Neural pattern plot is Fig. 3 also reveals that the ventral motor neurons (V*) have noticeably stronger pattern values than the dorsal motor neurons (D*), which implies that these neurons are responsible for a greater activity that creates the network oscillations.

In Fig. S4, we look at the pattern values for motor neurons to better identify the neurons responsible for the network behavior during forward motion. We consider a neuron "positive" or motivating forward motion if its pattern value is above the median and "negative" and less motivating if the pattern values are below the median. Using this method, we identify 32 active ventral motor neurons in each of the first two modes, 21 active dorsal motor neurons in the first mode, and 18 active dorsal motor neurons in the second mode.

In the first mode, we see 29 motor neurons that are positively activated (undulating upward), and 38 motor neurons that are negatively activated (undulating downward). In the second mode we find an opposite trend, with 37 motor neurons that are positively activated and 26 neurons that are negatively activated. Analysis of active neurons in each mode shows that there are a total of 74 active neurons, with 17 of them active only in one mode and 57 active in both modes. Notably, while such analysis can classify neurons into functional groups, it cannot directly represent the forces that will act on different segments of the body.

We also examine the first two time-dependent coefficients and see clear periodic oscillations in both. Summing up these two coefficients, we see the oscillations have a period of about 1.8 seconds and have roughly circular trajectory in the low-dimensional phase space. Such oscillations we observe in the time dependent coefficients support the correlation between the activity of the neural network and the body dynamics of the nematode; when *C. elegans* is swimming forward with its body following a periodic wave pattern, the activity of the neural network also tends to undergo periodic oscillations.

#### *VI.* VISCOELASTIC ROD MODEL

In order to represent *C. elegans* movement, we apply a discrete viscoelastic rod model that has been used to describe motion in anguilliform swimmers (18). The rod representation is chosen based on the anguilliform motion seen in *C. elegans* as it reacts to various mechanical touches and similar stimuli. In this model, the swimmer's body is represented as a two-dimensional rod composed of discrete rigid segments and joints, and forces acting on the rod represent the swimmer's muscle activity. The environment of the organism is modeled through damping, which may be varied to imitate different surrounding media (18,19). The model has similar principles to the spring based biomechanical model proposed for *C. elegans* (20). We chose to work with the former since it was derived through a consistent reduction of equations describing body in fluid into discrete segments, which shown to be numerically stable and efficient under dynamic neural stimulations. The model also includes an approximation of the environment that can describe various fluid media (e.g., agar, water) with configurable properties such as fluid viscosity. Notably, our results of body movement are generic, and we do not expect them to be essentially altered with various body models.

Here we summarize shortly the modeling approach and explain how we utilize viscoelastic rod model for *C. elegans* body. The state of the viscoelastic rod at any point in time is described by the x-y coordinates of the midpoint of each segment of the rod, and by the angle, $\varphi$, of each rod segment relative to the horizontal plane. We consider the joints connecting the segments of the rod to be actuated by passive springs, dashpots, and time-dependent force generators (as visualized in Fig. 1A). Given this model, force applied near one end of the rod can lead to movement that travels through the other end of the rod, depending on the magnitude of initial force and parameters governing the rod and its environment. The equations used to calculate the position of the segments of the viscoelastic rod are:

$$x_{i+1} = \frac{h}{2}\left(cos\ (\varphi_i) + cos\ (\varphi_{i+1})\right)\ + x_i \quad (6)$$

$$y_{i+1} = \frac{h}{2}\left(sin\ (\varphi_i) + sin\ (\varphi_{i+1})\right)\ + y_i \quad (7)$$

The differential equation determining the change in $\varphi$, based on the forces **f** and **g** applied to the segments of the rod in the x and y directions, respectively, is:

$$J\ddot{\varphi} = M_i - M_{i-1} + \frac{h}{2}(g_i + g_{i-1})\cos(\varphi) - \frac{h}{2}(f_i + f_{i-1})sin(\varphi) \quad (8)$$

The components of Eqn. (8) are defined as the contact moment $M_i$ (Eqn. (9)), the moment of inertia $J_i$ for link $i$ (Eqn. (10)), and the moment of inertia $I$ for motions in the x-y plane (Eqn. (11)), as:

$$M_i = EI_i\left(\frac{(\varphi_{i+1} - \varphi_i)}{h} - k_i\right) + \delta_i\left(\frac{(\dot{\varphi}_{i+1} - \dot{\varphi}_i)}{h}\right) \quad (9)$$

$$J_i = \ \rho h\left(I_i + \frac{\pi}{12}r^2h^2\right) \quad (10)$$

$$I = \frac{\pi D^4}{64} \quad (11)$$

In Eqn. (9), we solve for the contact moment based on the preferred curvature of the rod $k_i$, the rod's elasticity $E$ (Young's modulus), the environmental damping coefficient $\delta$, and the segment lengths $h_i$. The parameters of equations defining the moments of inertia $J$ and $I$ Eqn. (10, 11) depend on the rod's material density $\rho$, rod radius $r$, segment length $h_i$ and rod diameter $D$.

Expanding Eqn. (8) by substituting in Eqn. (9 - 11) we solve for $\ddot{\varphi}_i$ as follows:

$$J_i\ddot{\varphi} = M_i - M_{i-1} + \frac{h}{2}(g_i + g_{i-1})\cos(\varphi_i) - \frac{h}{2}(f_i + f_{i-1})sin(\varphi_i) \quad (12)$$

$$\rho h\left(I_i + \frac{\pi}{12}r^2h^2\right)\ddot{\varphi} = \ EI_i(\frac{(\varphi_{i+1} - \varphi_i)}{h} - k_i) + \delta_i(\frac{(\dot{\varphi}_{i+1} - \dot{\varphi}_i)}{h}) - EI_{i-1}(\frac{(\varphi_i - \varphi_{i-1})}{h} - k_i)$$
$$-\delta_{i-1}\left(\frac{(\dot{\varphi}_i - \dot{\varphi}_{i-1})}{h}\right) + \frac{h}{2}(g_i + g_{i-1})cos(\varphi_i) - \frac{h}{2}(f_i + f_{i-1})sin(\varphi_i) \quad (13)$$

$$\ddot{\varphi}_i = EI_i(\frac{(\varphi_{i+1} - \varphi_i)}{h} - k_i) + \delta_i(\frac{(\dot{\varphi}_{i+1} - \dot{\varphi}_i)}{h}) - EI_{i-1}(\frac{(\varphi_i - \varphi_{i-1})}{h} - k_i) - \delta_{i-1}(\frac{(\dot{\varphi}_i - \dot{\varphi}_{i-1})}{h})$$
$$+\frac{h}{2}(g_i + g_{i-1})cos(\varphi_i) - \frac{h}{2}(f_i + f_{i-1})sin(\varphi_i) \cdot \frac{1}{\rho h\left(I_i + \frac{\pi}{12}r^2h^2\right)} \quad (14)$$

We then transform Eqn. (14) into a system of first order differential equations:

$$p_1 = \varphi;\ p_2 = \dot{\varphi};\ \dot{p}_1 = \dot{\varphi} = p_2 \quad (15)$$

$$\dot{p}_{2i} = \ddot{\varphi}_i = \{EI_i\left(\frac{\left(p_{1(i+1)} - p_{1i}\right)}{h} - k_i\right) + \delta_i\left(\frac{\left(p_{2(i+1)} - p_{2i}\right)}{h}\right)$$
$$-EI_{i-1}\left(\frac{\left(p_{1i} - p_{1(i-1)}\right)}{h} - k_i\right) - \delta_{i-1}\left(\frac{\left(p_{2i} - p_{2(i-1)}\right)}{h}\right) + \frac{h}{2}(g_i + g_{i-1})\cos(p_{1i})$$
$$-\frac{h}{2}(f_i + f_{i-1})\sin(p_{1i}) \cdot \frac{1}{\rho h\left(I_i + \frac{\pi}{12}r^2h^2\right)} \quad (16)$$

which can be solved computationally. We then include calcium dynamics defined in (18),

$$\ddot{\beta} + c_1\dot{\beta} + c_2\beta = c_3u(t) \quad (17)$$
$$\ddot{\eta} + \dot{\eta}c_4 + c_5\eta = c_6\beta(t) \quad (18)$$
$$A(t) = \frac{a_0 + (\rho\eta)^2}{1 + (\rho\eta)^2} \quad (19)$$

where *u(t)* is each motor neuron output in terms of $\beta$, the T-tubuli depolarization response and $\eta$, the SR calcium release, with constant parameters $c_1 = 60$, $c_2 = 20$, $c_3 = 50$, $c_4 = 10$, $c_5 = 30$, $c_6 = 30$. SR calcium release $\eta$ is used to define *A(t)*, the activation function with $\rho(t) = 1$, representing muscle fatigue.

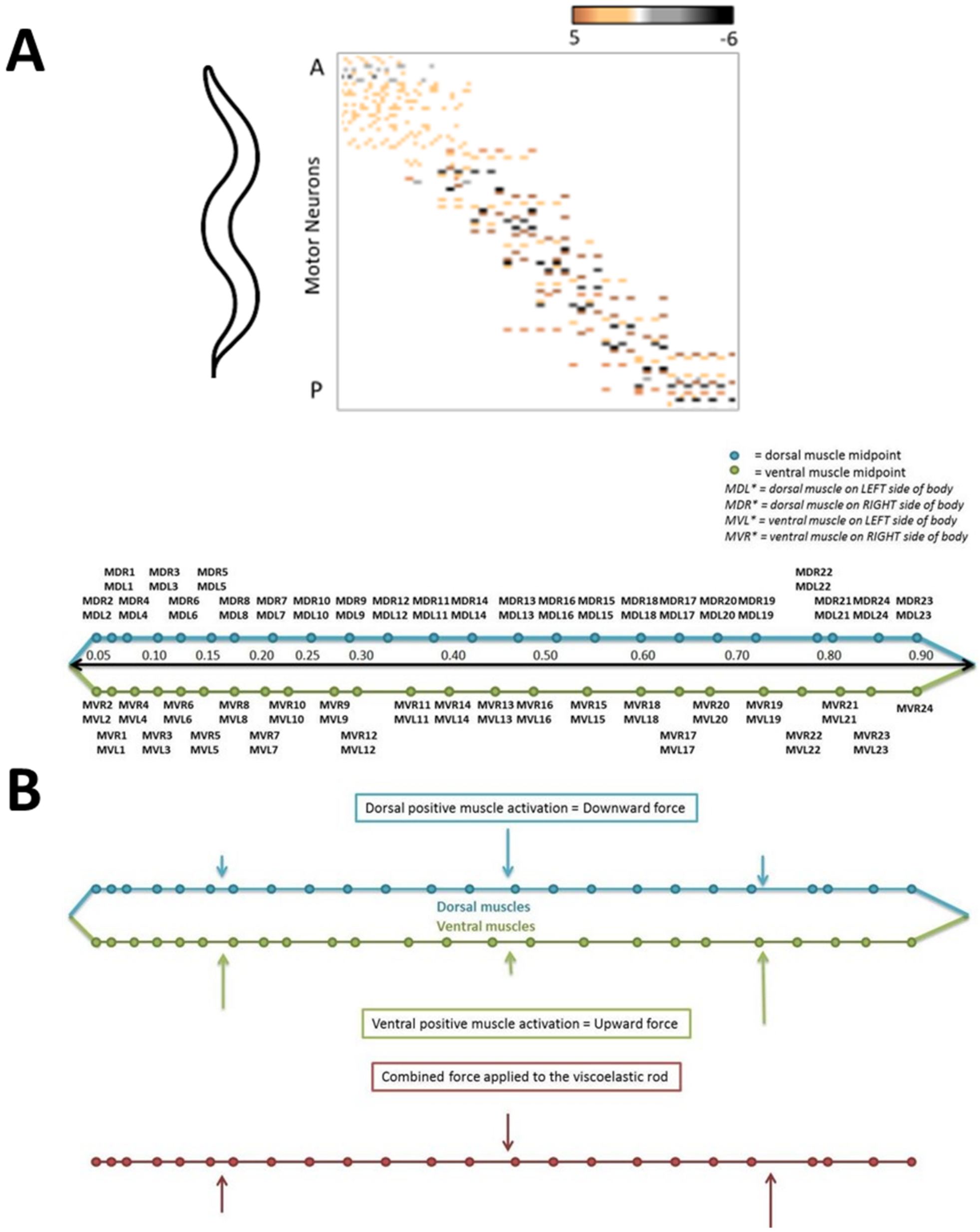


**Figure S5: A:** Top: Connectivity from motor neurons to muscles ordered from anterior (top) to posterior (bottom) direction. Bottom: Map of the 95 muscles represented as segments of the viscoelastic rod. Segment length is determined by position of each muscle relative to the AP-axis, with the overall body length scaled to 1. **B**: Force is applied to segments of the viscoelastic rod to represent muscle activation. Dorsal and ventral muscles are combined based on their locations relative to the AP-axis and summing the input into each group of muscles determines the overall force applied to a segment of the rod.

#### *VII.* VISCOELASTIC ROD AS A MODEL FOR *C. ELEGANS* BODY

We apply the viscoelastic rod model to *C. elegans* by representing each muscle group in the nematode as a segment of the rod connected with joints. We approximate the forces applied to the rod segments using neuron activity data from forward motion simulations and a neuron-to-muscle map which are experimentally determined (21). The combined activity of the neurons connected to a muscle is considered the muscle's input, which leads to activation of force.

To represent accurate *C. elegans* musculature, we model 95 muscles relevant to *C. elegans* locomotion and use their approximate sizes and locations to determine segment length and position. These 95 individual muscles are divided into four groups based on their physical location in the nematode: dorsal left (DL), dorsal right (DR), ventral left (VL), and ventral right (VR). Every muscle is assigned a position along the Anterior-Posterior axis (AP-axis) of the nematode, and the dorsal left and dorsal right muscles are coupled into single segments representing the dorsal muscle groups, and likewise for the ventral muscles. Dorsal (ventral) left muscles are paired only with dorsal (ventral) right muscles where left-right pairs are determined from muscle location along the AP-axis, as shown in Fig. S5A.

Differentiation between the dorsal and ventral muscle groups is necessary since these two muscle groups act in opposition to each other; for forward motion to occur, the dorsal muscles contract while the ventral muscles relax, and vice versa. As such, we model activation of the muscles in the dorsal group through downward force, and the muscles in the ventral group through upward force. We then merge the dorsal and ventral muscle groups based on their location along the AP-axis to form a single discrete rod. With such representation, the dorsal (downward) and ventral (upward) forces can be summed over the dorsal-ventral muscle pairs to determine the overall force applied to each segment of the rod: for example, if a large downward force is applied by a dorsal muscle and a small upward force is applied by the corresponding ventral muscle, the net result will be application of downward force on the segment of the rod representing the merged muscles (Fig. S5B).

Unlike the generic anguilliform swimmer model, the basic *C. elegans* viscoelastic rod representation assumes force is only applied in the y-direction, meaning **f** is a zero vector. Given the absence of force in the x-direction, we eliminate the sine term from Eqn. (18), resulting in the final system of equations:

$$\dot{p}_{1i} = \dot{\varphi}_i = p_{2i} \tag{20}$$

$$\dot{p}_{2i} = \ddot{\varphi}_i = EI_i\left(\frac{\left(p_{1(i+1)} - p_{1i}\right)}{h} - k_i\right) + \delta_i(\frac{\left(p_{2(i+1)} - p_{2i}\right)}{h}) - EI_{i-1}\left(\frac{\left(p_{1i} - p_{1(i-1)}\right)}{h} - k_i\right) - \delta_{i-1}\left(\frac{\left(p_{2i} - p_{2(i-1)}\right)}{h}\right) + \frac{h}{2}(g_i + g_{i-1})\cos(p_{1i}) \cdot \frac{1}{\rho h\left(I_i + \frac{\pi}{12}r^2h^2\right)} \tag{21}$$

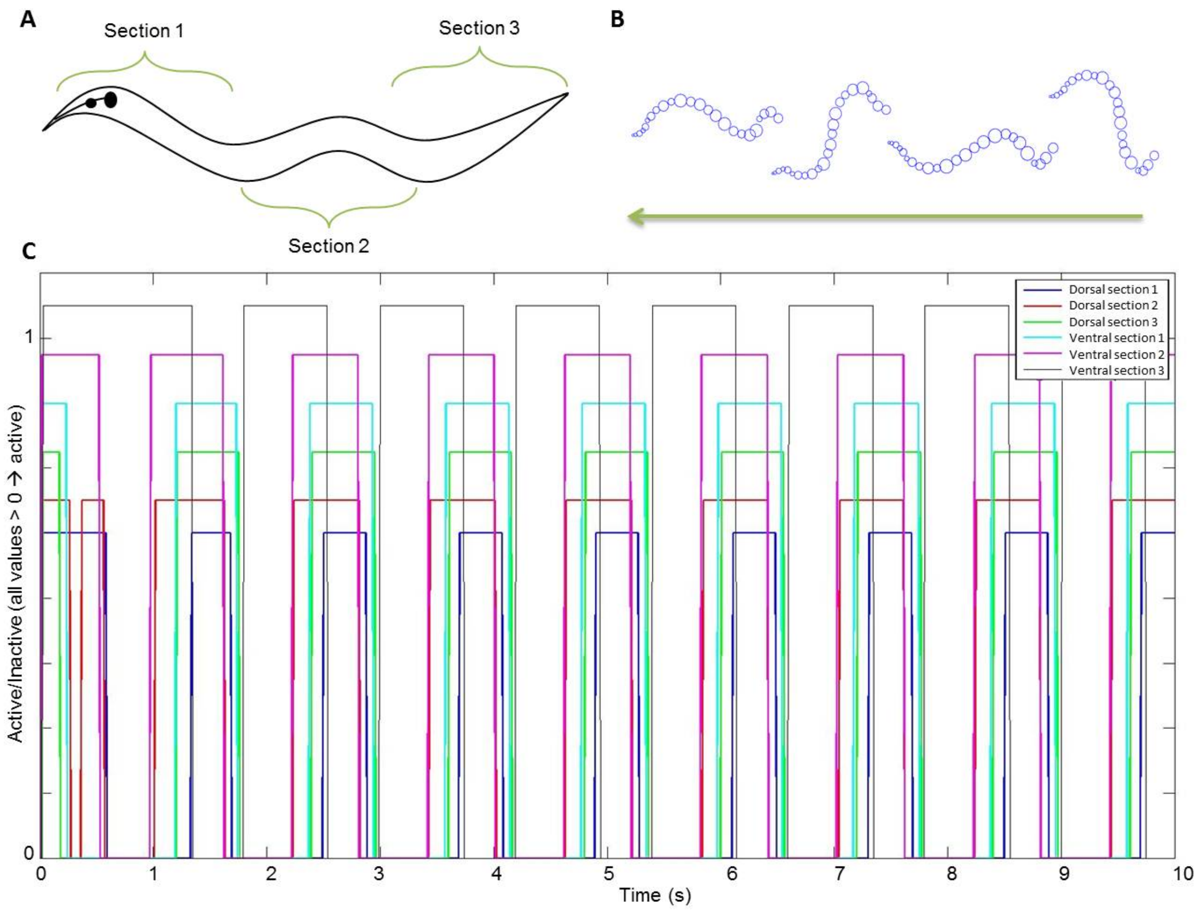


**Figure S6: A**: Three main contraction/relaxation sections. It is expected that dorsal and ventral muscles will not continuously be activated to contract simultaneously and that the stimulation of the three sections of the worm will be staggered **B**: Results of the simulations of viscoelastic rod movement over time during FWD stimulation. The worm is moving from left to right, with the size of each rod segment represented by the diameter of each circle. **C**: Activation patterns of the groups of dorsal/ventral muscles based on input from connected motor neurons during FWD excitation showing the repeating activation pattern of the six muscle groups.

for modeling *C. elegans* as a viscoelastic rod.

The parameters defining the remaining physical characteristics of the rod were set based on published analyses of high-speed images of *C. elegans* forward-swimming in a controlled environment (22) and micropipette deflection experimentation on anesthetized *C. elegans* (23). We model an adult *C. elegans* with diameter D=65μm, Young's modulus E=3.77-8 kPA and material density ρ=1.0 $g/(cm^3)$. We assume the preferred curvature to be the equilibrium state of the nematode, represented by the zero vector, and assign damping coefficient δ = 1 Ns/m as an approximate representation of the substrates making *up C. elegans* habitat (broadly ranging from rotting vegetation to animal intestines). As previously described, we assign segment lengths to vector $h_i$ based on approximate muscle length. These body-environmental parameters are fully configurable through modWorm.

The musculature and body model dynamics are computed from the normalized membrane voltages of motor neurons (i.e. $\overline{V_i} = V_i - V_{th}$). Since the system described by viscoelastic rod model is non-stiff, it can be solved using explicit integration method. We use RK4 explicit ODE solver for Python and DP5 solver for Julia. **Computational time is optimized to match real-time with a ratio of 1:1**, i.e., 1 sec of integrated body dynamics ~ 1 sec of actual body dynamics. Notably, the simulation time step for body is set to match that of nervous system dynamics, thus allowing simulation of both neural and body dynamics in the same time sampling frequency. Like neural simulations, multiple body simulations can be run in parallel utilizing modWorm's ensemble simulation function.

The behavior of the viscoelastic body model is validated by the known contraction and relaxation patterns found in the nematode's body during forward motion. These patterns closely align with the simulated formations of the viscoelastic rod as it moves over time. The behavior of the viscoelastic rod as it models step input currents for FWD, BWD and TURN scenarios shows a clear relationship between stimulation of motor neurons along the body and muscle movement (Fig. S7). Since the model is sensitive to the magnitude of motor neuron stimulation a muscle receives, we observe that greater force is applied in certain sections of the worm over time, creating the undulating form associated with *C. elegans* forward motion. Specifically, we often see the anterior and posterior sections receiving force in the opposite direction of the force applied to the middle section of the worm (Fig. S6B). For example, FWD motion configuration represents dorsal (ventral) contraction in the anterior and posterior sections with ventral (dorsal) contraction in the middle section, which is consistent to the expected muscle usage.

##### *C. ELEGANS* BODY SHAPES DURING MOTION

| Time | FWD | BWD | TURN |
|---|---|---|---|
| 0.05 | 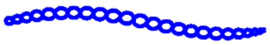 | 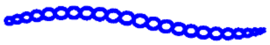 | 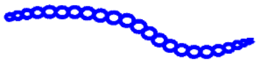 |
| 0.15 | 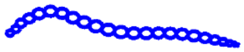 | 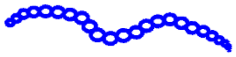 | 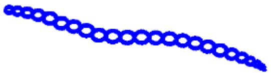 |
| 0.25 | 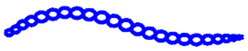 | 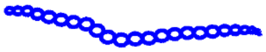 | 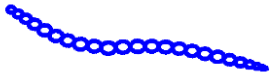 |

| | | | |
|---|---|---|---|
| 0.35 | 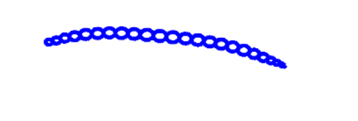 | 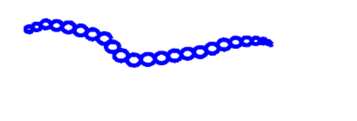 | 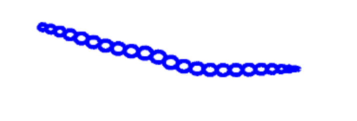 |
| 0.45 | 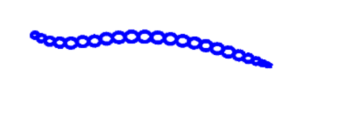 | 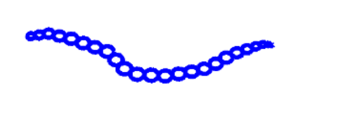 | 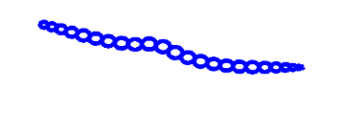 |
| 0.55 | 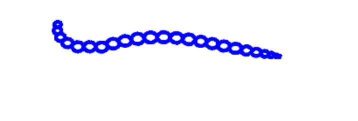 | 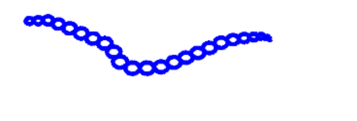 | 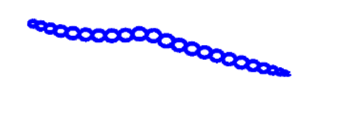 |
| 0.65 | 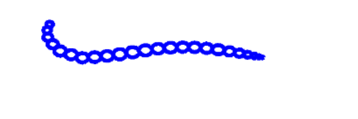 | 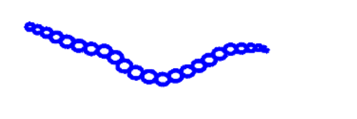 | 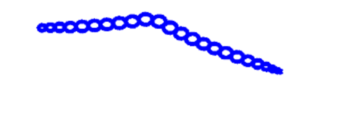 |
| 0.75 | 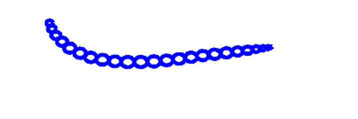 | 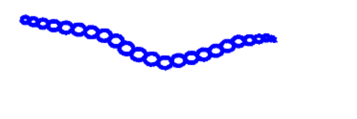 | 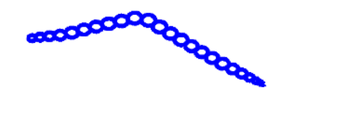 |
| 0.85 | 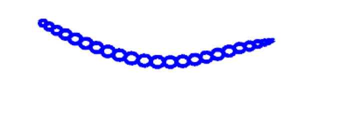 | 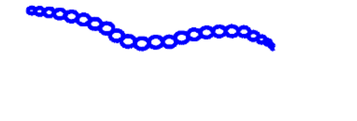 | 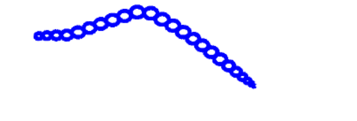 |
| 0.95 | 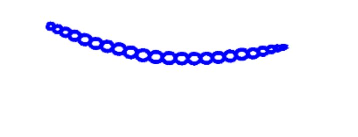 | 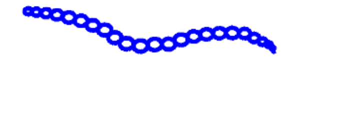 | 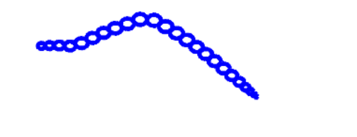 |
| 1.05 | 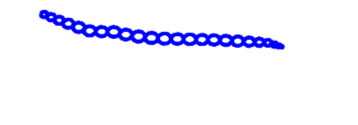 | 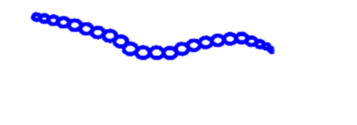 | 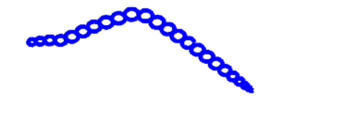 |
| 1.15 | 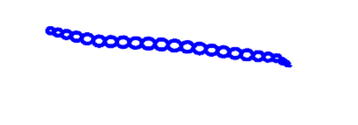 | 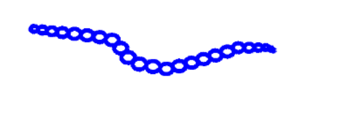 | 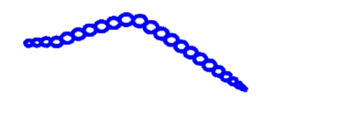 |
| 1.25 | 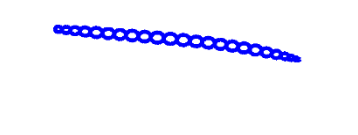 | 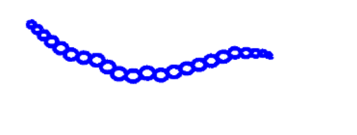 | 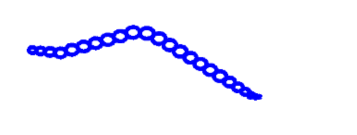 |
| 1.35 | 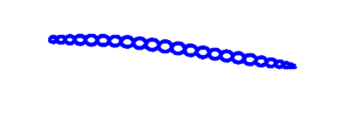 | 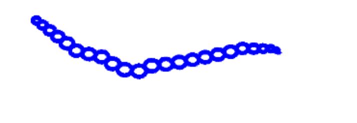 | 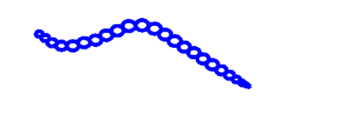 |
| 1.45 | 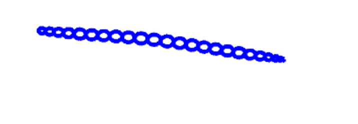 | 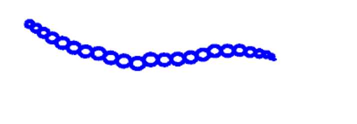 | 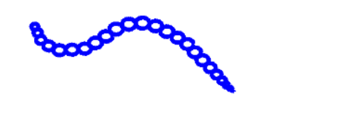 |
| 1.55 | 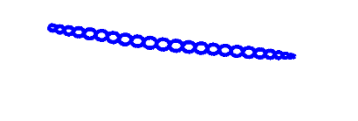 | 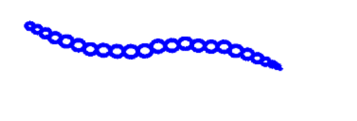 | 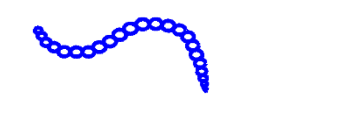 |

| | | | |
|---|---|---|---|
| 1.65 | 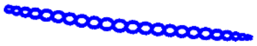 | 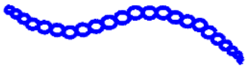 | 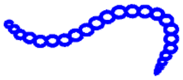 |
| 1.75 | 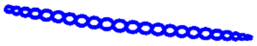 | 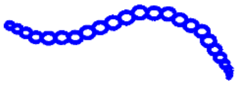 | 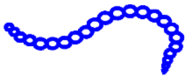 |
| 1.85 | 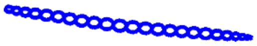 | 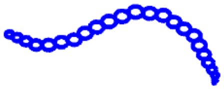 | 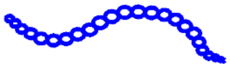 |
| 1.95 | 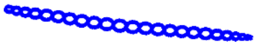 | 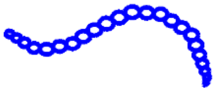 | 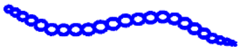 |
| 2.05 | 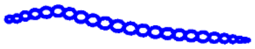 | 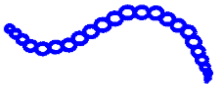 | 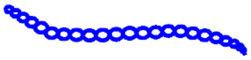 |

**Figure S7:** Library of body shapes seen during simulation of forward, backward, and turning motion induced by constant neural stimuli. Backward and turning shapes are more extreme than the smooth shapes associated with forward motion.

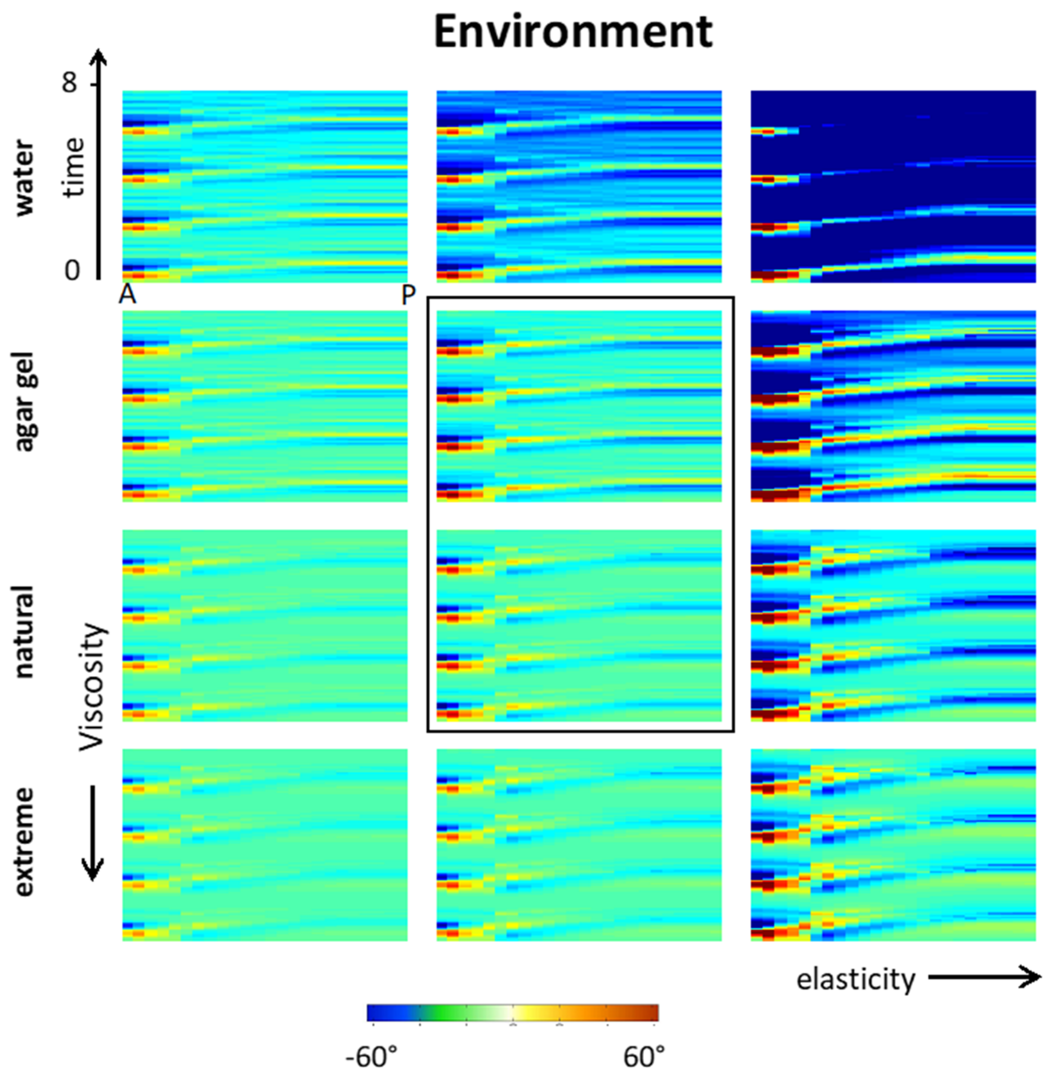


**Figure S8:** Effect of viscosity variation and body elasticity on forward movement invoked by constant stimulus (viscosity: 1, $10^2, 10^3, 10^4$ $cP$; elasticity: 3.75, 6,8. 25 kPA). Only specific choice of parameters permits coherent movement.

#### *VIII.* ENVIRONMENTAL VARIATIONS

Experiments indicate that the environment plays a significant role in shaping coordinated movement (24,25). In Fig. S8, we vary fluid viscosity and body elasticity to study their effects on simulated body postures (i.e. curvature plots) during forward locomotion as characterized in Fig. 4, S7. Increasing viscosity values impedes movement by breaking off wave propagation from anterior to posterior, whereas decreasing them causes rapid extreme strokes unlike the body shapes seen in efficient *C. elegans* forward motion. When elasticity variations are added, these strokes intensify and create atypical movements for high elasticity or no movement for low elasticity.

#### *IX.* BACKWARD INTEGRATION FROM MUSCLE FORCE TO NEURAL VOLTAGE

We add backward integration to our model to approximate muscles to neural dynamics interaction in addition to neurons stimulating muscles (26,27). The procedure is as follows: for an arbitrary external force acting on muscles at time *t*, the force can be approximated in the form of activation signals to muscles described as column vectors $\vec{E}_D(t), \vec{E}_V(t)$ each with dimension $(24 * 1)$ for dorsal and ventral segments of the body respectively. These vectors are then expanded into $\vec{E}_D^{Left}(t), \vec{E}_D^{Right}(t), \vec{E}_V^{Left}(t), \vec{E}_V^{Right}(t)$ where:

$$\vec{E}_D^{Left}(t) = \vec{E}_D^{Right}(t) = \vec{E}_D(t) * \frac{1}{2} \tag{22}$$

$$\vec{E}_V^{Left}(t) = \vec{E}_V^{Right}(t) = \vec{E}_V(t) * \frac{1}{2} \tag{23}$$

These 4 vectors account for the full profile of forces throughout the dorsal, ventral, left and right axis of the body where left and right forces are assumed to be symmetric in the direction of left and right axis of the body. $\vec{E}_D^{Left}(t), \vec{E}_D^{Right}(t), \vec{E}_V^{Left}(t), \vec{E}_V^{Right}(t)$ are then vertically stacked to a single column vector $\vec{E}_{DV}(t)$ of dimension $(96 * 1)$, which describes the muscle forces for the whole body of the worm at time t.

We then solve the inverse problem of forward integration process (i.e. neural dynamics to muscle forces). During the forward integration, muscle forces at time $t$ is the linear mapping of neural voltages $V(t)$ (279 * 1) via muscle mapping matrix $M(96 * 279)$:

$$\vec{E}_{D,V}(t) = MV(t) \tag{24}$$

Matrix $M$is of *fat* type (i.e. more columns than rows), meaning that for a given force $\vec{E}_{D,V}(t)$, there exist multiple neural voltage solutions which satisfy Eqn. 24. But since $M$ is not of full rank, we cannot solve $V(t)$ with ordinary least squares method. This can be circumvented using generalized pseudo-inverse conceived by Singular Value Decomposition. Using this technique, we approximate the neural voltages from muscle forces by solving the following inverse equation.

$$\vec{V}_E(t) = M^{+}\vec{E}_{D,V}(\mathrm{t}) \tag{25}$$

Here, $M^{+} = \mathrm{Vs}^{-1}\mathrm{U}^{\mathrm{T}}$ is the generalized pseudo-inverse of $M$ computed via taking the inverse of Singular Value Decomposition of $M$ where U and V correspond to unitary matrices each containing left and right singular vectors of M and s is the diagonal matrix whose diagonal entries are the singular values of M. Notably, Eqn. 25 effectively solves for the least norm solution $\vec{V}_E(t)$ that satisfies

$$\left|M\vec{V}(t) - \vec{E}_{D,V}(t)\right|_2 \geq \left|M\,\vec{V}_E(t) - \vec{E}_{D,V}(t)\right|_2 \tag{26}$$

i.e., $\vec{V}_E(t)$ is the least norm solution which minimizes the Euclidean norm of error (28).

The approximated voltage $\vec{V}_E(t)$ can be interpreted as the inferred neural voltage at time *t* induced by muscle forces $\vec{E}_{D,V}(t)$. To drive the nervous system with $\vec{V}_E(t)$, we define the superposed voltage:

$$V_i^{sp} = V_i + V_{E_i} \tag{27}$$

where $V_i^{sp}$ is the voltage state of ith neuron superposed with $V_{f_i}$. Equivalently, Eqn. 27 can be written as a current term induced by muscle force $I_i^{force}(\vec{\boldsymbol{V}}_{E_i})$. The modified equation allows driving the nervous system entirely by muscle to neural dynamics interactions. Specifically, the network dynamics driven by $\vec{V}_E$ is given by:

$$C\dot{V_i}(t) = -G^C\left(V_i^{sp} - E_{cell}\right) - I_i^{Gap}\left(\vec{\boldsymbol{V}}^{sp}\right) - I_i^{Syn}\left(\vec{\boldsymbol{V}}^{sp}\right) \tag{28}$$

where

$$I_i^{Gap} = \sum_j G_{ij}^{g}\left(V_i^{sp} - V_j^{sp}\right) \tag{29}$$

and

$$I_i^{Syn} = \sum_j G_{ij}^{s} s_j\left(V_i^{sp} - E_j\right) \tag{30}$$

$$s_i = a_r\phi(V_i;\beta,V_{th})(1 - s_i) - a_d s_i \tag{31}$$

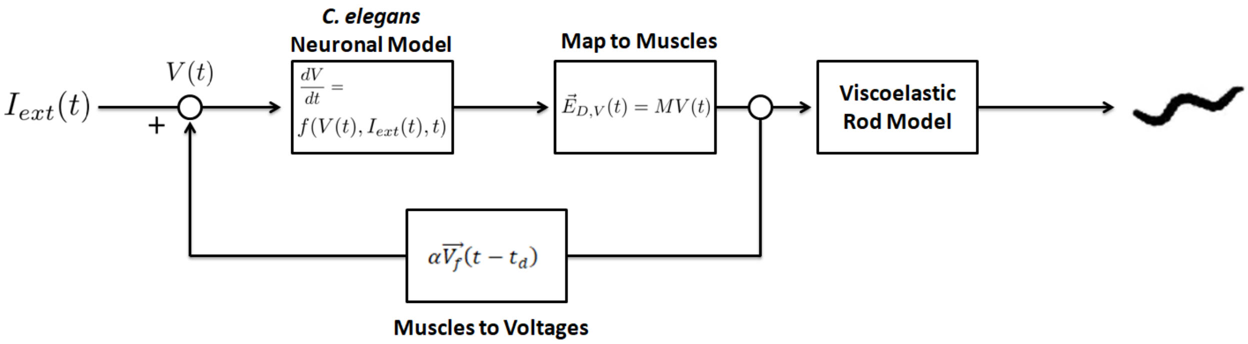


**Figure S9:** Block diagram of proprioceptive feedback implementation. $I_{ext}(t)$ is the input stimuli into nervous system at time *t*, $V(t)$ is the neuron voltages at time *t*, $\vec{E}_{D,V}$(t) is muscle forces at time *t*, and M is the muscle mapping matrix.

$$\phi(v_i; \beta, V_{th}) = \frac{1}{1 + \exp\left(-\beta\left(V_i^{sp} - V_{th}\right)\right)} \tag{32}$$

The solution $V(t)$ then describes the neural activity driven by $\vec{V_E}$ which satisfy the right-hand side of Eqn. 28. Notably, Eqn. (28 - 32) are nonlinear and therefore computed neural voltage at $t + \Delta t$ transformed to muscle force is not guaranteed to be identical to the initial arbitrary muscle force $(\vec{E_D}(t), \vec{E_V}(t))$. Such selectivity of force by the nervous system is best described in Fig. 3. The difference between initial force and output force by the nervous system can also occur when the external neural stimulation occurs simultaneously with external muscle force stimulations.

#### *X.* PROPRIOCEPTIVE FEEDBACK

We use muscle forces to neurons inverse integration to emulate proprioceptive feedback from the environment. We assume that when the muscles exert forces on the environment, there are reactive forces from the particles in the local environment that are proportional to the muscle forces. Such interaction is particularly evident in distinct locomotion patterns of *C. elegans* in different fluids such as agar vs water (25). We model the environment-body interaction through an approximation in which environmental reactive forces are sensed by the nervous system with a particular delay. We implement the delayed feedback as follows: at each time $t$ muscle forces are transformed to voltages $\vec{V_E}(t)$ according to Eqn. (25). Once simulation has passed certain time $t > t_{init}$, e.g., duration of initial pulse of input stimuli associated with a mechanical touch, the voltages $\vec{V}(t)$ in Eqn. (1 - 5) is set to be superposition of intrinsic voltages $\vec{V}$ and voltages $\vec{V_E}$ induced by delayed proprioceptive feedback:

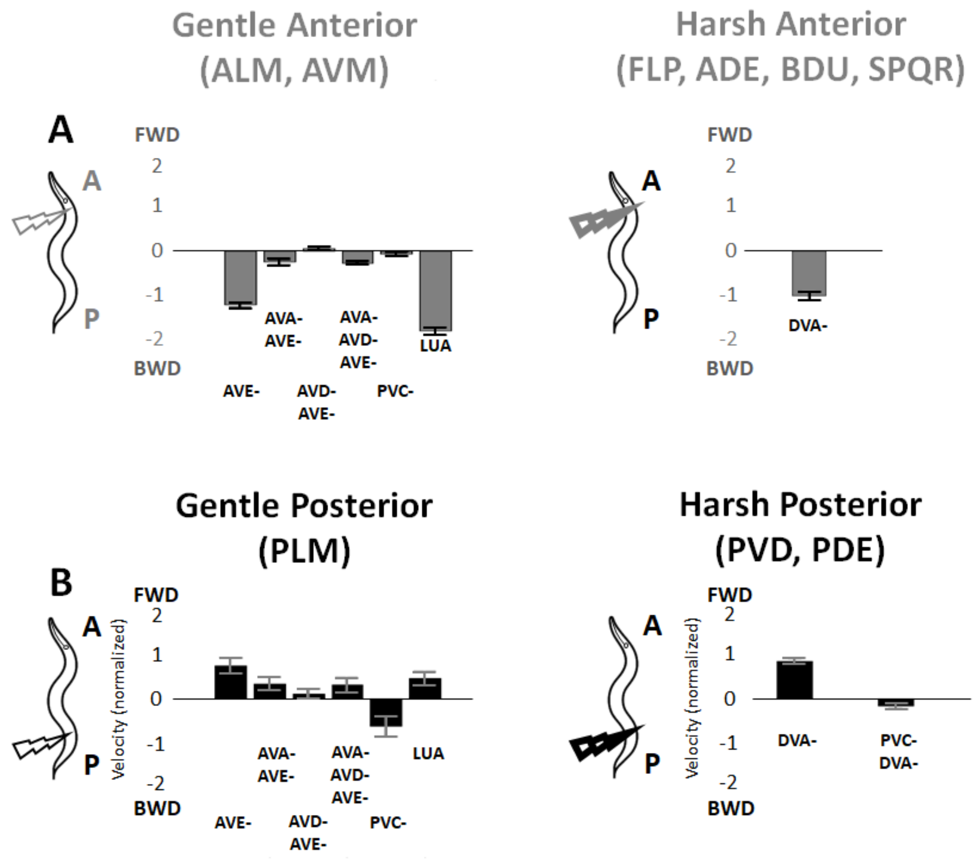


**Figure S10**: Additional *in-silico* ablations for gentle and harsh touch responses. **A:** Additional ablations for anterior gentle and harsh touch responses. **B:** Additional ablations for posterior gentle and harsh touch responses (Supplement to Figure 6).

$$\vec{V}(t) \rightarrow \vec{V}(t) + \alpha\overrightarrow{V_E}(t - t_d) \quad (33)$$

where $\alpha$ is the scaling factor and $t_d$ is the time delay. Equivalently, this is identical to adding a current term $I_i^{force}(\vec{\boldsymbol{V}}_{E_i})$ to the Eqn. 1. The block diagram of the implementation is shown in Fig. S9.

We show in Fig. 4 the effect of the delay on the curvature and find optimal values. In Fig. S11, we show motor neurons voltage profiles for movements enabled by feedback such as forward, backward locomotion and composite movements such as avoidance. Movements are triggered by short pulse of neural stimuli, which are gradually turned off exponentially, and sustained by feedback.

#### *XI.* OPTOGENETIC ABLATION OF AVA

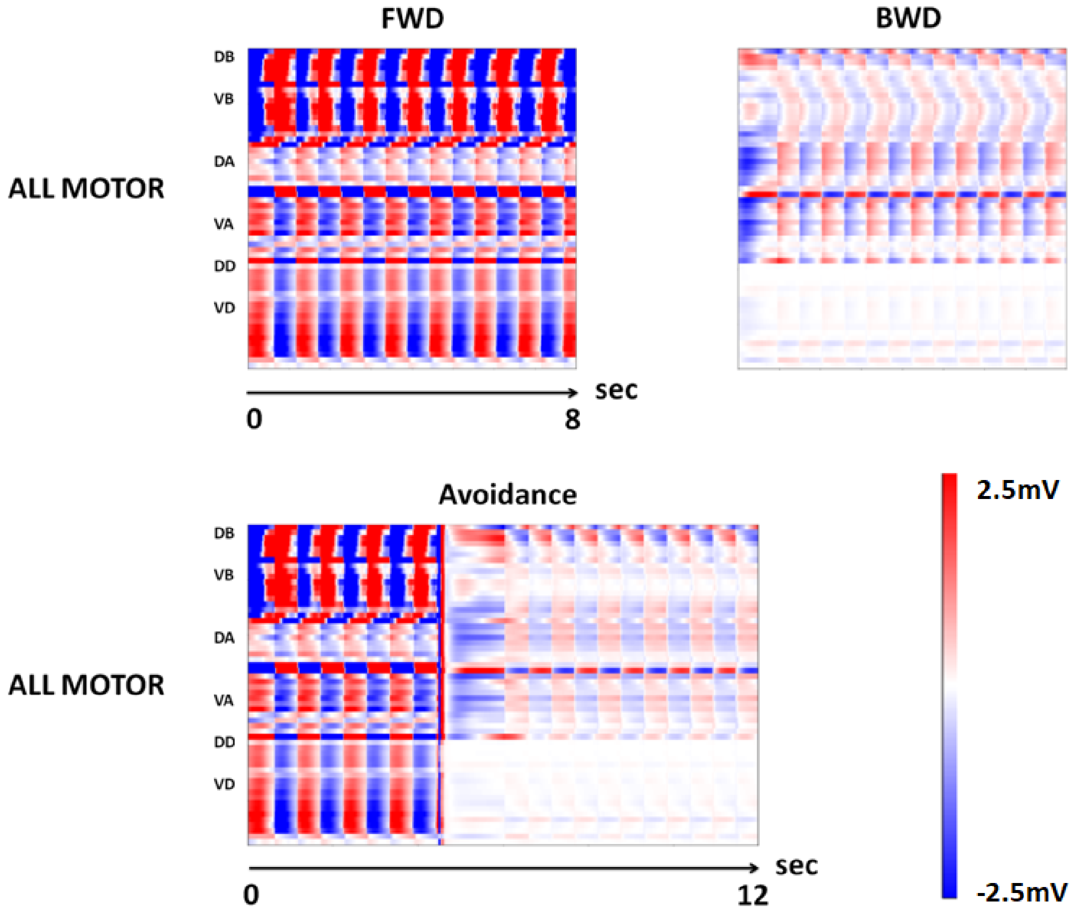


**Figure S11**: Motor neurons dynamics for forward, backward, and avoidance behaviors with proprioceptive feedback (supplement to Figure 7).

Standard culture methods were used to maintain strains (29). Animals were grown on nematode growth medium (NGM) plates and maintained on a diet of the *E. Coli* strain OP50. The N2 Bristol strain was used as the wild-type control. The strain ZM7198 (*lin-15(n765ts); hpEx3072*[*Prig-3*-MiniSOG::SL2::wCherry] +pL15EK[*lin-15AB* genomic DNA]) was used for AVA ablation (30). All strains were maintained at a low population density in dark conditions.

**AVA ablation.** We used the strain ZM7198 expressing the miniature singlet oxygen generator (miniSOG) in AVA neurons under control of the *rig-3* promoter and exposed animals to blue light to kill neurons through the production of reactive oxygen species, as previously described (31). Briefly, 20-30 larval stage 4 (L4) animals were washed in M9 buffer to remove residual food and placed on an unseeded NGM plate surrounded by a ring of 150 mM $CuSO_4$ to prevent worms from crawling away during illumination. Blue light was delivered for 12 minutes using a 488 nm 50 W LED (Mightex Systems) at a 50% duty cycle with a pulse width of 0.25 seconds using a Pulser USB light train generator (Prizmatix,

Ltd.). The plate was positioned at approximately 15 cm from the light source such that the intensity of light delivered was 200 mW/cm$^2$ as measured by a PM100D optical power meter (Thorlabs, Inc.). Animals were allowed to recover for at least 1 hour in the dark before behavioral assays. AVA ablation was verified by uncoordinated backing in response to anterior touch (32). For control experiments wild-type N2 animals were subjected to the same light stimulation protocol described above.

***In-vivo* behavioral assays.** Animals that had been allowed to recover after illumination were transferred to 6 cm NGM plates with a thin bacterial lawn. Fresh plates were made daily by spreading 30 µl of OP50 evenly across the surface of the plate and incubated at room temperature overnight. Touch assays were performed as previously described (32). Briefly a sable hair from a paint brush was taped to a glass pipette and used to gently touch animals near the tail. The video was recorded for up to a minute following touch or until the animal left the field of view. Behavioral data were aligned relative to the time of touch for comparison. Video recordings were made on a Nikon SMZ800 trinocular microscope (Nikon Instruments, Inc.) with a Pike F421b digital camera (Allied Vision Technologies, GmbH) using Fire-I 6.0 image acquisition software (Unibrain, Inc.) at a rate of 15 frames per second. Videos were analyzed using WormLab 3.1 worm tracking software (MBF bioscience) to obtain velocity, bending amplitude and position data for individual animals. Graphs were generated using Matlab R2019a (The Mathworks, Inc.) and Graphpad Prism 8.2.1 (GraphPad Software, Inc.).

#### *XII.* COMPUTING EIGENWORM COEFFICIENTS FROM SIMULATIONS

Recalling from section VI, the viscoelastic model solves the segment angles $\varphi_i$ ($i \in [1,..,24]$) at each timepoint alongside the body in respect to the horizontal line in x-y plane. Since eigenworm coefficients assume there are 48 such angles along the body, we extrapolate $\varphi_i$ to span $i \in [1,..,48]$. After extrapolation, we normalize $\varphi_i$ in respect to mean angle $\overline{\varphi_i}$ for each timestep and project experimentally obtained eigenworm modes via performing dot product between $\varphi_t$ and $M$ where $M$ is (48 * 7) matrix where the columns consist of first 7 normalized eigenworm modes. Once we have projections for all timepoints, we then obtain the coefficients $s_i$ for first 7 eigenworm modes by computing the 2-norm for each projected mode (i.e. column) alongside the time axis. Finally, we obtain the normalized coefficients through $s_i / \sum s_i$. The normalized coefficients error with respect to the empirical values is then computed by calculating the mean absolute error between simulated vs experimentally obtained normalized coefficients as follows:

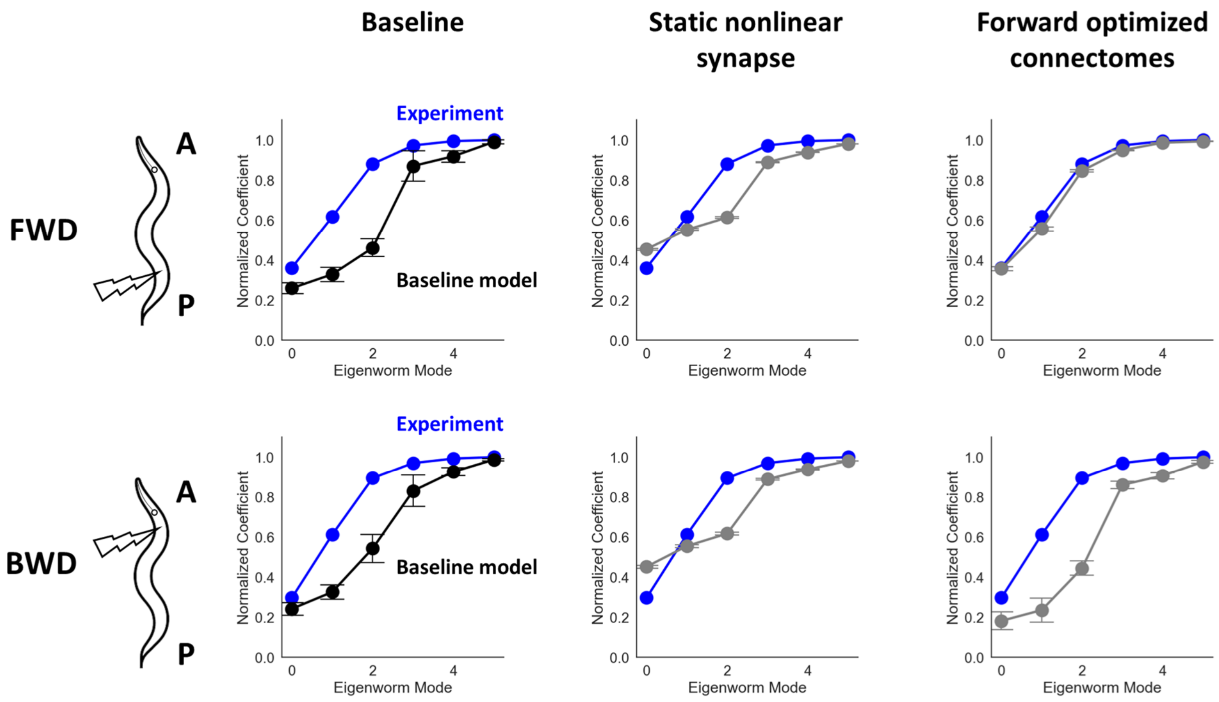


**Figure S12**: The cumulative eigenworm coefficients for forward and backward locomotion with respect to experimental coefficients (blue). From left to right, cumulative coefficients for the first 6 modes using baseline mode, baseline model using static nonlinear synapses of the form $s \sim \sigma(v)$ instead of equation 4 and using optimized connectomes with respect to forward locomotion coefficients.

$$error = \frac{1}{n}\sum_{i=1}^{n}|s_i - \hat{s}_i| \tag{34}$$

#### *XIII.* CONNECTOME OPTIMIZATION

We use the genetic algorithm to optimize the synapse strengths of connectome data provided by (33) where we optimize the scaling factors $\alpha_{ij}$ to each existing synapse weight $w_{ij}$ for both gap and synaptic connectomes, amounting to total 5146 scaling factors to be optimized. The scaling factor range is set to [0.5, 1.5] and the Mean Absolute Error (MAE) formula defined by Eqn. 34 is used as an objective function. For the hyperparameters of Genetic Algorithm we use the population size of 17 with 100 generations with crossover probability of 95% and mutation probability of 70%.

#### *XIV.* STATIC NONLINEAR SYNAPSES

We replace the original synaptic transmission model described by Eqn. (3, 4) with simpler static nonlinearity model using sigmoid function as follows:

$$s = \sigma(V) \tag{35}$$

$$\sigma(V) = \frac{1}{1 + e^{-\beta * V}} \tag{36}$$

Where $V$ is the neuron membrane potential and $\beta$ is the sigmoid width parameter and set to $-0.125$ identical to the value used in Eqn. 5.

#### *XV.* ONLINE BLOG FOR EXPLORATIONS OF BEHAVIORAL SCENARIOS

We started a blog to share and elaborate on various stimulation and behavioral scenarios we have studied in the paper (Fig. S14). The blog will also host additional scenarios that will enhance the model and potentially contribute to better understanding of mapping neural dynamics to body movements. Through the blog we also aim to interact with the community and get it involved in identifying novel scenarios.

The blog is available as part of the Github repository:
**https://shlizee.github.io/modWorm/**

#### A Initialize nervous system class

```
class CelegansWorm_NervousSystem:

    def __init__(self):
```

Network size

```
        self.network_Size = n_params.CE.n
```

Neurons' properties

```
        self.neuron_C = n_dyn.init_neuron_C(capacitance = n_params.CE.cell_caps) # (n,)

        self.neuron_Linear = n_dyn.init_neuron_Linear(conductance = n_params.CE.leak_conductances,
                                                      leak_voltage = n_params.CE.leak_potentials) # (2 x n)

        self.neuron_Chemical = n_dyn.init_neuron_Chemical(synaptic_rise_time = n_params.CE.synaptic_rise_tau,
                                                          synaptic_fall_time = n_params.CE.synaptic_fall_tau,
                                                          sigmoid_width = n_params.CE.B) # (3 x n)
```

Connections + interactions properties

```
        self.network_Electrical = n_inter.init_network_Electrical(conn_map = n_params.CE.gap_conn_2011_haspel,
                                                                  conductance_map = n_params.CE.gap_conductances,
                                                                  active_mask = np.ones(279, dtype = 'bool')) # (n x n)

        self.network_Chemical = n_inter.init_network_Chemical(conn_map = n_params.CE.syn_conn_2011_haspel,
                                                              conductance_map = n_params.CE.syn_conductances,
                                                              polarity_map = n_params.CE.ei_map,
                                                              active_mask = np.ones(279, dtype = 'bool')) # (2 x n x n)
```

#### B Using new connectome mapping

New adjacency matrices for Electrical and Chemical connectomes

```
# Initialize network and its properties
self.network_Electrical = n_inter.init_network_Electrical(conn_map = gap_conn_Cook_2019,
                                                          conductance_map = n_params.CE.gap_conductances,
                                                          active_mask = np.ones(279, dtype = 'bool')) # (n x n)

self.network_Chemical = n_inter.init_network_Chemical(conn_map = syn_conn_Cook_2019,
                                                      conductance_map = n_params.CE.syn_conductances,
                                                      polarity_map = n_params.CE.ei_map,
                                                      active_mask = np.ones(279, dtype = 'bool')) # (2 x n x n)
```

#### C Incorporating non-linear neural channels (AWA)

```
self.neuron_EGL19_AWA = n_dyn.init_neuron_Nonlinear(self, channel_type = 'EGL19_awa',
                                                    neuron_inds = [73, 82],
                                                    params_mat = EGL19_params,
                                                    added_order = 1,
                                                    initconds_mat = EGL19_initcond,
                                                    using_julia = True)

self.neuron_SHK1_AWA = n_dyn.init_neuron_Nonlinear(self, channel_type = 'SHK1_awa',
                                                   neuron_inds = [73, 82],
                                                   params_mat = SHK1_params,
                                                   added_order = 2,
                                                   initconds_mat = SHK1_initcond,
                                                   using_julia = True)
```

#### D Incorporating tyramine gated chloride channels (LGC-55)

```
self.network_Chemical = n_inter.init_network_Chemical(conn_map = n_params.CE.syn_conn_2019_nw_haspel,
                                                      conductance_map = n_params.CE.syn_conductances,
                                                      polarity_map = ei_map_LGC55,
                                                      active_mask = np.ones(279, dtype = 'bool')) # (2 x n x n)
```

New adjacency matrix describing excitatory-inhibitory mapping

**Figure S13**: Using modWorm interface to incorporate each variation to the model. **A:** Defining a nervous system class prior to the simulation. "init_" functions allow defining each nervous system module with user defined parameters. **B:** To initialize the model with non-baseline connectome data (e.g. dataset from Cook et al, 2019), simply provide NumPy arrays (279 * 279) each corresponding to desired electrical and chemical connectome mappings. **C:** To initialize the model with neurons with non-linear channels, define self.channel_name on top of baseline model to incorporate neural channels to particular set of neurons with their respective parameters. **D:** To initialize the model with modified synaptic polarities, simply replace the "ei_map" parameter of init_network_Chemical() function with new NumPy array (279 * 279) incorporating modified neuron to neuron polarities (e.g., LGC-55). (supplement to Figure 7).

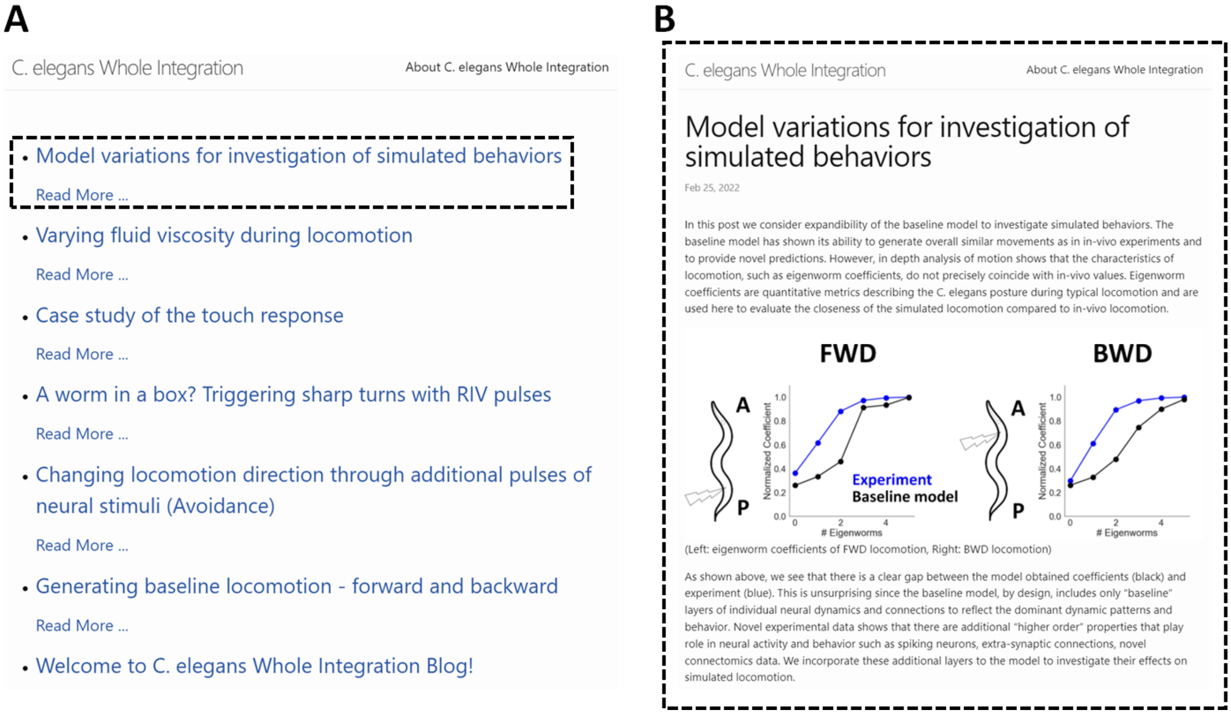


**Figure S14:** Online blog hosting the scenarios in Figure 2-7 and future additional studies. **A:** The main page of the blog featuring the studied scenarios. **B:** Redirected page after clicking on "Read More" on "Model variations for investigation of simulated behaviors" post.
